# A Hypothalamic Circuit Underlying the Dynamic Control of Social Homeostasis

**DOI:** 10.1101/2023.05.19.540391

**Authors:** Ding Liu, Mostafizur Rahman, Autumn Johnson, Iku Tsutsui-Kimura, Nicolai Pena, Mustafa Talay, Brandon L. Logeman, Samantha Finkbeiner, Seungwon Choi, Athena Capo-Battaglia, Ishmail Abdus-Saboor, David D. Ginty, Naoshige Uchida, Mitsuko Watabe-Uchida, Catherine Dulac

## Abstract

Social grouping increases survival in many species, including humans^1,2^. By contrast, social isolation generates an aversive state (loneliness) that motivates social seeking and heightens social interaction upon reunion^3-5^. The observed rebound in social interaction triggered by isolation suggests a homeostatic process underlying the control of social drive, similar to that observed for physiological needs such as hunger, thirst or sleep^3,6^. In this study, we assessed social responses in multiple mouse strains and identified the FVB/NJ line as exquisitely sensitive to social isolation. Using FVB/NJ mice, we uncovered two previously uncharacterized neuronal populations in the hypothalamic preoptic nucleus that are activated during social isolation and social rebound and that orchestrate the behavior display of social need and social satiety, respectively. We identified direct connectivity between these two populations of opposite function and with brain areas associated with social behavior, emotional state, reward, and physiological needs, and showed that animals require touch to assess the presence of others and fulfill their social need, thus revealing a brain-wide neural system underlying social homeostasis. These findings offer mechanistic insight into the nature and function of circuits controlling instinctive social need and for the understanding of healthy and diseased brain states associated with social context.

## Main Text

Throughout geographical and cultural settings, humans readily form groups and flourish in social activities^7^. People with strong social networks tend to be happier, healthier, and better in coping with stress, while prolonged deprivation of social interaction is associated with harmful effects including increased anxiety, fragmented sleep, impaired cognition, weakened immune system, and increased risk for cardiovascular illnesses and cancer^7,8^. Similarly, in the wild, animal grouping decreases the risk of predation, reduces energy consumption, and lends itself to mutually beneficial behaviors such as parenting and group foraging^1,2^. By contrast, experiments performed in a variety of species showed that social isolation leads to abnormal behaviors and brain activity, and increased disease susceptibility^5,9,10^. The aversive state of social isolation may thus act as an evolutionarily conserved alarm signal in individuals separated from a group to enhance sensitivity to social cues and promote social seeking behavior^6,8^. Strikingly, the lack of social drive is a hallmark of several neurodevelopmental and psychiatric disorders^8,11^, such as autism, depression, and schizophrenia, although a neural underpinning of this impairment remains elusive.

Essential physiological needs such as those for food, water or sleep are encoded by brain circuits that monitor the organism’s metabolic, osmotic and sleep/wake state in order to signal emerging needs — hunger, thirst or drowsiness — and motivate the corresponding goal-directed survival behaviors — eating, drinking or sleeping. The intensity of the restoring behavior matches the level of deprivation, leading to satiation when the organism’s homeostasis is reached^12^. The hypothalamus has emerged as a brain hub underlying physiological homeostasis. Specifically, various hypothalamic neuronal populations play key roles in orchestrating distinct survival needs. Examples include neurons in the arcuate nucleus expressing *AgRP* and *POMC* in the control of food appetite and satiety, respectively^13,14^, neurons in lamina terminalis and MnPO in water intake^15-17^, and various hypothalamic populations in sleep control^18^. Intriguingly, experiments in rodents show that increased durations of social isolation trigger increasingly stronger rebounds in social interaction^3,4^. Thus, the need for social interaction may follow a homeostatic process driven by the discomfort (“loneliness”) elicited by social deprivation in a similar manner to other physiological needs^6^.

What neural mechanisms underlie the need for social interaction? Recent studies have identified the role of dopamine-, oxytocin- and serotonin-associated brain circuits in mediating social motivation and reward upon reunion^19-22^. However, the brain response to isolation and its impact on circuits driving social interaction remain elusive. Here we hypothesized that, in addition to the previously identified nodes underlying social reward, specific hypothalamic circuits may exist that underlie social homeostasis.

Work described here uncovers the identity and function of distinct genetically defined populations of neurons in the mouse hypothalamus that encode social need and social satiety. Circuit tracing and functional analyses identify associated brain-wide neural networks underlying the homeostatic control of social grouping and point to the sense of touch as the sensory modality that informs animals about their social context. Our findings provide new insights into the neural basis of instinctive social drive and may provide new avenues to the understanding of social behavior in normal and pathological contexts.

## Social rebound as a behavioral manifestation of social homeostasis

Laboratory mice prefer grouped over isolated housing^19,23^, providing an attractive experimental system to investigate the neural underpinnings of social need. Sibling cage mates (4-5 weeks) were separated and singly housed for up to 5 days, and the subsequent rebound in social interaction during reunion was quantified as a proxy for the underlying social need (Fig. 1a, b). Experiments were performed in adult female mice to avoid interfering behaviors such as aggression or mating seen in males. We examined 6 distinct mouse strains to assess their respective levels of social drive. Most strains tested displayed significant rebounds in social interaction after isolation (Fig. 1c-e), suggesting that social isolation enhances social need and promotes social activity. The strength of social rebound, measured as total interaction time, interaction bout number and bout duration, steadily increased while the distance between animals (social distance) and the behavioral latency decreased with longer durations of isolation (Fig. 1c-e and Extended Data Fig. 1a-c), providing a direct measure of the increased social need elicited by lengthier isolation. Intriguingly, different strains displayed highly diverse ranges of social rebound, from absent or weak (BALB/c and DBA), to moderate (C57BL/6J, C3H/HeJ, SWR/J), and strong (FVB/NJ), suggesting various sensitivities to social isolation according to the genetic background (Fig. 1c-e and Extended Data Fig. 1a-c). Importantly, the strength of rebound in each strain was not correlated with the reported stress sensitivity for that strain^24,25^. Based on these data, we selected the FVB strain for further behavioral and functional experiments.

**Figure 1.**
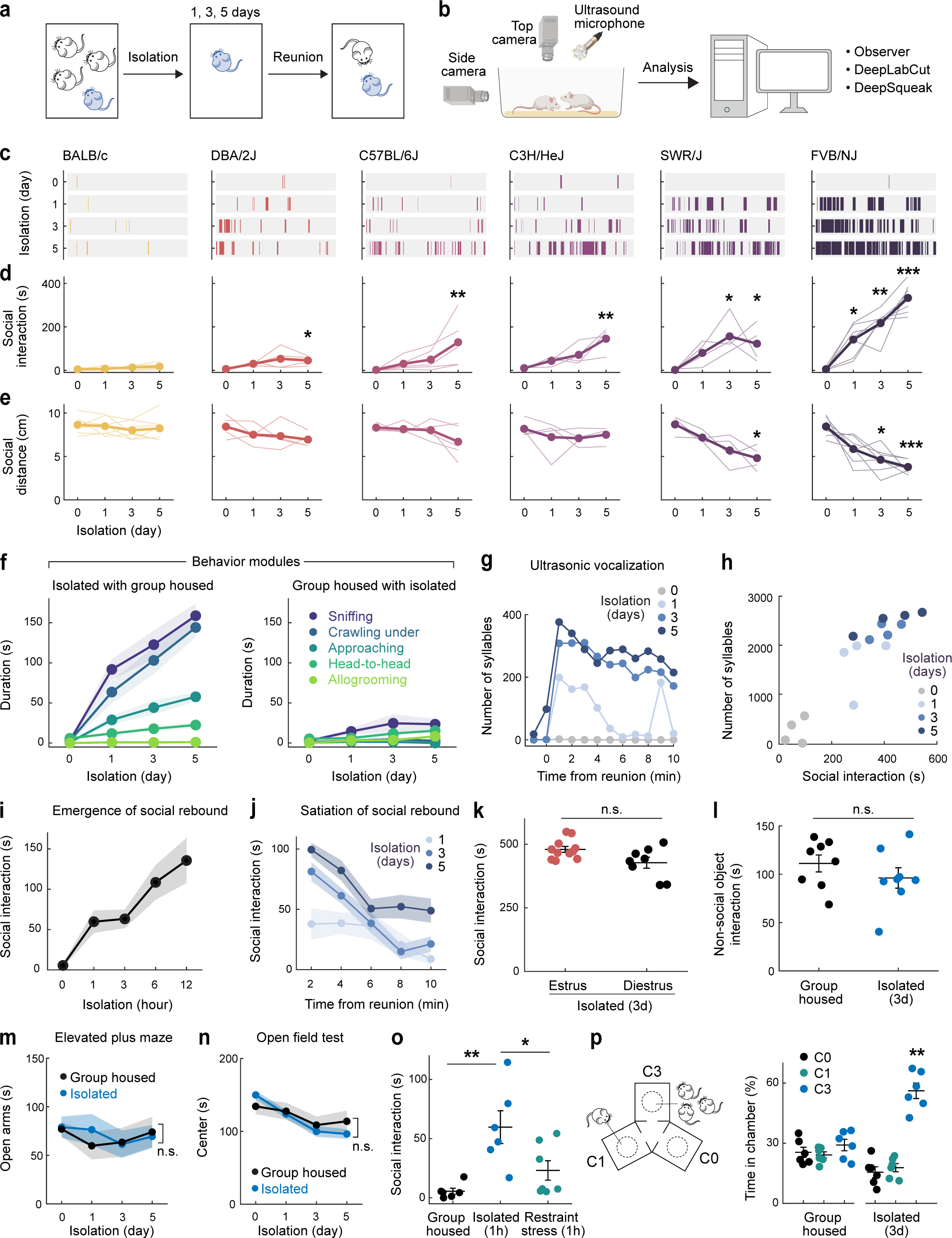
Social rebound as behavior manifestation of social homeostasis. **a, b** Short-term social isolation/reunion behavioral paradigm and analysis pipeline. **c**, Raster plots of social events during reunion on example mouse from each strain. **d, e** Total duration of social interaction and average distance between two mice (social distance) during 10 min social reunion. Thin lines represent individual mice, thick lines are cohort average (BALB/c n=7, DBA/2J n=4, C57BL/6J n=5, C3H/HeJ n=4, SWR/J n=4, FVB/NJ n=7). **f**, Behavioral modules displayed during social rebound by previously isolated (left) versus paired group housed FVB mice (right), n=7 pairs. **g**, Ultrasonic vocalization (USV) during social reunion from example FVB mouse. **h**, Correlation between duration of social interaction and number of USV syllables during social reunion. Each dot represents a mouse, n=4. **i**, Emergence of social rebound in FVB mice following increasing time in isolation, n=6. **j**, Satiation of social rebound after various lengths of isolation, n=7. **k**, Social rebound in different phases of the estrous cycle, estrus phase n=11, diestrus phase n=8. **l**, Investigation of non-social object in group housed versus isolated FVB mice, n=8. **m, n**, Elevated plus maze test (n=5) and open field test (n=9) in isolated vs group housed FVB mice. **o**, Social interaction in FVB mice after 1h of social isolation, n=6 or body restraint stress, n=7. **p**, Preference test among empty (C0), one-mouse (C1) and 3-mice chambers (C3), n=6. **d, e**, Friedman test between baseline (day 0) and each isolation day; **k, l, o**, Mann–Whitney U test; **m, n**, Two-way ANOVA; **p**, Friedman test; n.s., not significant; *p<0.05, **p<0.01, ***p<0.001. All shaded areas and error bars represent the mean ± s.e.m.

Social rebound episodes were subdivided into distinct behavioral modules such as approaching, sniffing, crawling under, head-to-head contact and allogrooming (Fig. 1f, Extended Data Fig. 1d, e and Supplementary Video 1) and were associated with robust ultrasonic vocalizations (Fig. 1g, h and Extended Data Fig. 1f-h). Although these sequences entailed reciprocal interactions between two mice placed together, behavior events were mostly initiated by the previously isolated mouse (Fig. 1f and Extended Data Fig. 1e). Behavior analysis following shorter isolation durations (1-12h) enabled us to characterize the emergence of social rebound with progressively increased intensity (Fig. 1i). By contrast, the display of rebound behavior declined over time during reunion, indicating a gradual satiation of social drive through social interaction (Fig. 1j). In control experiments, we showed that social rebound was independent from the female estrus stage (Fig. 1k), was not associated with a general increase in investigation behavior toward an object (Fig. 1l) or in stress responses (Fig. 1m-o) and that mice of different ages showed similar social rebound (Extended Data Fig. 1i). Interestingly, isolated mice preferred interacting with a group rather than a single mouse (Fig. 1p) and, in contrast to group housed mice, showed no preference for novel over familiar mice (Extended Data Fig. 1j, k), supporting the idea that social rebound is an affiliative behavior that compels animals back to the group.

FVB mice suffer from progressive retinal degeneration due to their homozygosity of the *Pde6b^rd1^* allele^26^. To assess the effect of visual impairment on rebound behavior, we tested the *Pde6b^rd1^/Pde6b^rd1^* mutation in the C57 strain background. We observed no difference in the rebound behavior of C57 *Pde6b^rd1^/Pde6b^rd1^* compared to C57 wildtype mice (Extended Data Fig. 1l). Moreover, *Pde6b^rd1/+^* offspring of FVB X C57 crosses which, unlike *Pde6b^rd1^/Pde6b^rd1^* mice, have functional vision^27^ exhibited significantly higher social rebound than the C57 strain (Extended Data Fig. 1m), suggesting that the elevated social rebound seen in FVB mice results from unique features in the genetic background of the FVB strain that are distinct from their impaired vision.

Together, these results show that short-term social isolation/reunion assays in FVB females offer a robust and naturalistic paradigm to investigate the neurobiological mechanisms underlying social need.

## Identification of candidate neurons underlying social homeostasis

The hypothalamus is a control hub for physiological homeostasis, such that activation of specific hypothalamic populations drives distinct motivated behaviors associated with physiological needs and their satiation^13-18^. The observed rebound in social interaction following social deprivation suggests that similar homeostatic circuits may exist in the hypothalamus to control social need and satiety. To explore this hypothesis, we performed microendoscopy calcium imaging of neuronal activity in the medial preoptic nucleus (MPN), a hypothalamic nucleus involved in the control of social behavior^28,29^. Pan-neuronal MPN imaging in freely interacting mice undergoing social isolation and subsequent reunion (Fig. 2a, b) reveals unique neuronal activation patterns in which non-overlapping neuronal populations are activated during either isolation or reunion episodes (Fig. 2c, d, Extended Data Fig. 2a, b and Supplementary Video 2). Strikingly, neuronal ensembles active during social isolation were promptly inhibited upon social reunion and then re-activated upon removal of the social partner (Fig. 2c, d, Cluster #3). This population, referred to as MPN^Isolation^ neurons (26 ± 4% of responsive neurons), appears to track the animal’s isolation state, thus representing a promising candidate population to signal social need. In addition, we observed populations of neurons activated during reunion with distinct temporal dynamics (Fig. 2c, d, Cluster #1 and #2), referred to as MPN^Reunion^ neurons (74 ± 4% of responsive neurons), which may convey signals associated with social reunion. Principal component analysis (PCA) of ensemble activity across all simultaneously imaged neurons revealed a trajectory with an instantaneous change of representation upon reunion after isolation, and a slow shift back to an isolation state after removal of the social partner (Fig. 2e, f).

**Figure 2.**
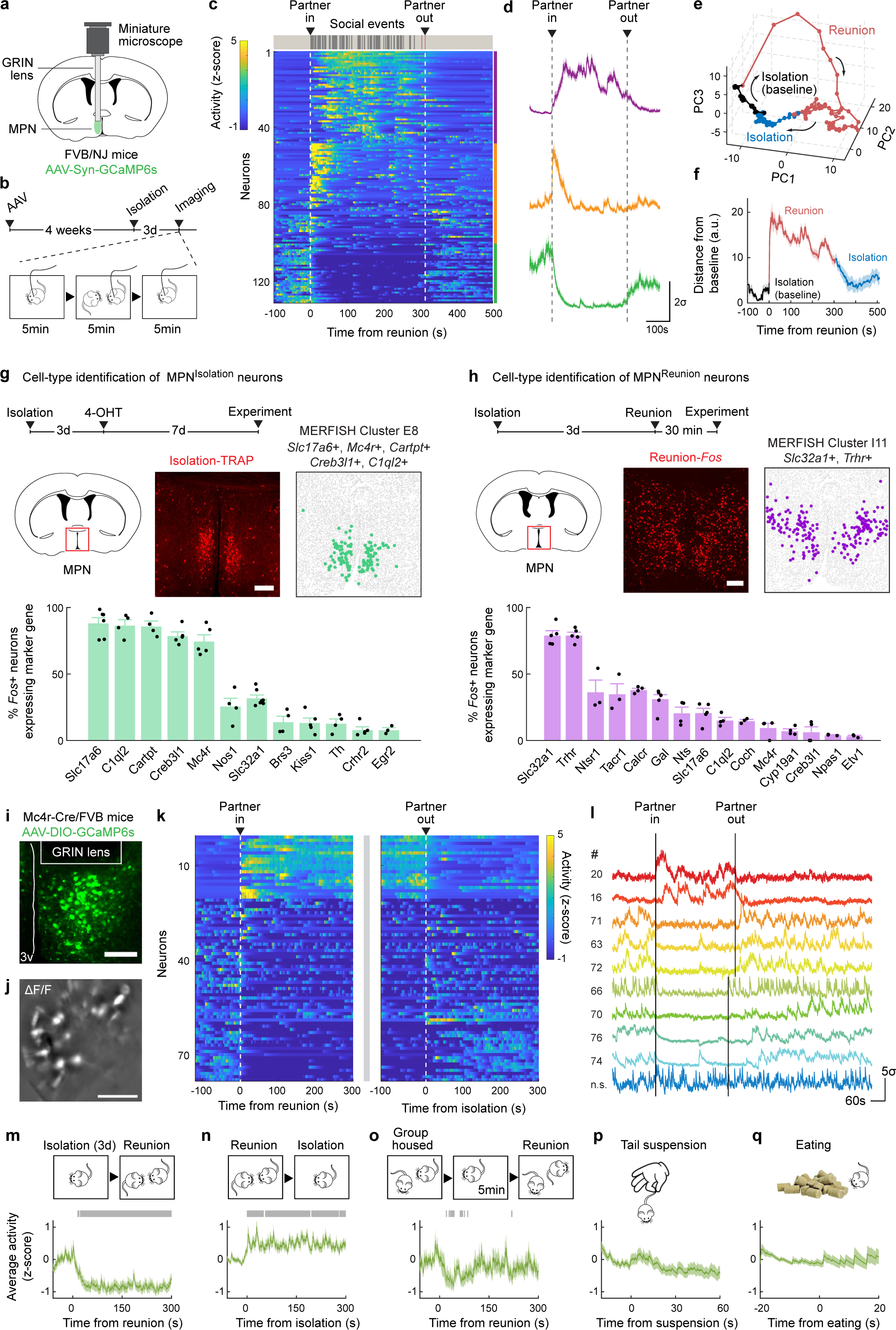
Identification of neuron types underlying social homeostasis. **a**, Microendoscopy calcium imaging in the hypothalamic medial preoptic nucleus (MPN). **b**, Behavioral and calcium imaging paradigm. **c**, MPN pan-neuronal activity during social isolation and reunion. Heatmap of activity in example mouse, n=3 mice. **d**, Average activity of neuronal populations from **c** with distinct activity patterns during isolation and reunion. **e**, Principal component analysis of population activity in **c**. Bin size = 5s. **f**, Distance from each PCA time point to the mean of baseline. Bin size = 1s. **g**,**h**, Cell type identification of MPN^Isolation^ (**g**) and MPN^Reunion^ (**h**) neurons based on overlap between activity induced *Fos* and expression of neuron-type marker genes, n=3-6 for each marker gene. **i**, Representative image showing MPN*^Mc4r^*^+^ neurons expressing GCaMP6s. **j**, Example imaging field of GRIN lens, represented as ΔF/F. **k**, MPN*^Mc4r^*^+^ neuronal activity during social isolation and social reunion. n=78 significantly modulated neurons pooled from 3 mice. **l**, Calcium activity of 10 example MPN*^Mc4r^*^+^ neurons. **m-q**, Average calcium activity of reunion-inhibited MPN*^Mc4r^*^+^ neurons in distinct behavioral regimes. Gray bars indicate significance of activity above or below 95% confidence interval of baseline. All shaded areas and error bars represent the mean ± s.e.m. unless otherwise noted. All scale bars, 200μm.

To molecularly identify candidate hypothalamic populations underlying the control of social need and satiety, we took advantage of the induction of the immediate early gene *Fos* as a readout of neuronal activity following behavior. Specifically, we used the activity-based and tamoxifen-inducible Cre line TRAP2^17^ crossed to the reporter line Ai9 to label integrated activity across several hours during social isolation (Fig. 2g) as well as *Fos in situ* hybridization to visualize transiently activated neurons during social rebound (Fig. 2h). Importantly, isolation-activated neurons in the MPN were not activated by group housing, restraint stress or food deprivation (Extended Data Fig. 2c). To precisely identify MPN isolation and reunion neurons, we sought to assign these populations to our recently established spatial transcriptomic cell-type atlas in the preoptic region^29^. Using *in situ* hybridization, we examined the overlap between *Fos* labeled neurons and 20 selected marker genes identifying 14 transcriptomic MPN neuronal populations (Extended Data Fig. 3). The best fit for the population of MPN^Isolation^ neurons was the published MERFISH cluster E8^29^, a glutamatergic (*Slc17a6*+) population enriched in *Mc4r*, *Cartpt*, *Creb3l1* and *C1ql2* expression (Fig. 2g and Extended Data Fig. 3, 4a). In turn, the majority (∼80%) of MPN^Reunion^ neurons were GABAergic (*Slc32a1*+) and enriched in *Trhr* expression, thus corresponding to the previously described MERFISH cluster I11 (Fig. 2h and Extended Data Fig. 3, 4b). These results successfully assigned specific transcriptomically defined cell types to isolation- and reunion-activated neuronal populations in MPN and thus enabled direct genetic access for functional characterization.

**Figure 3.**
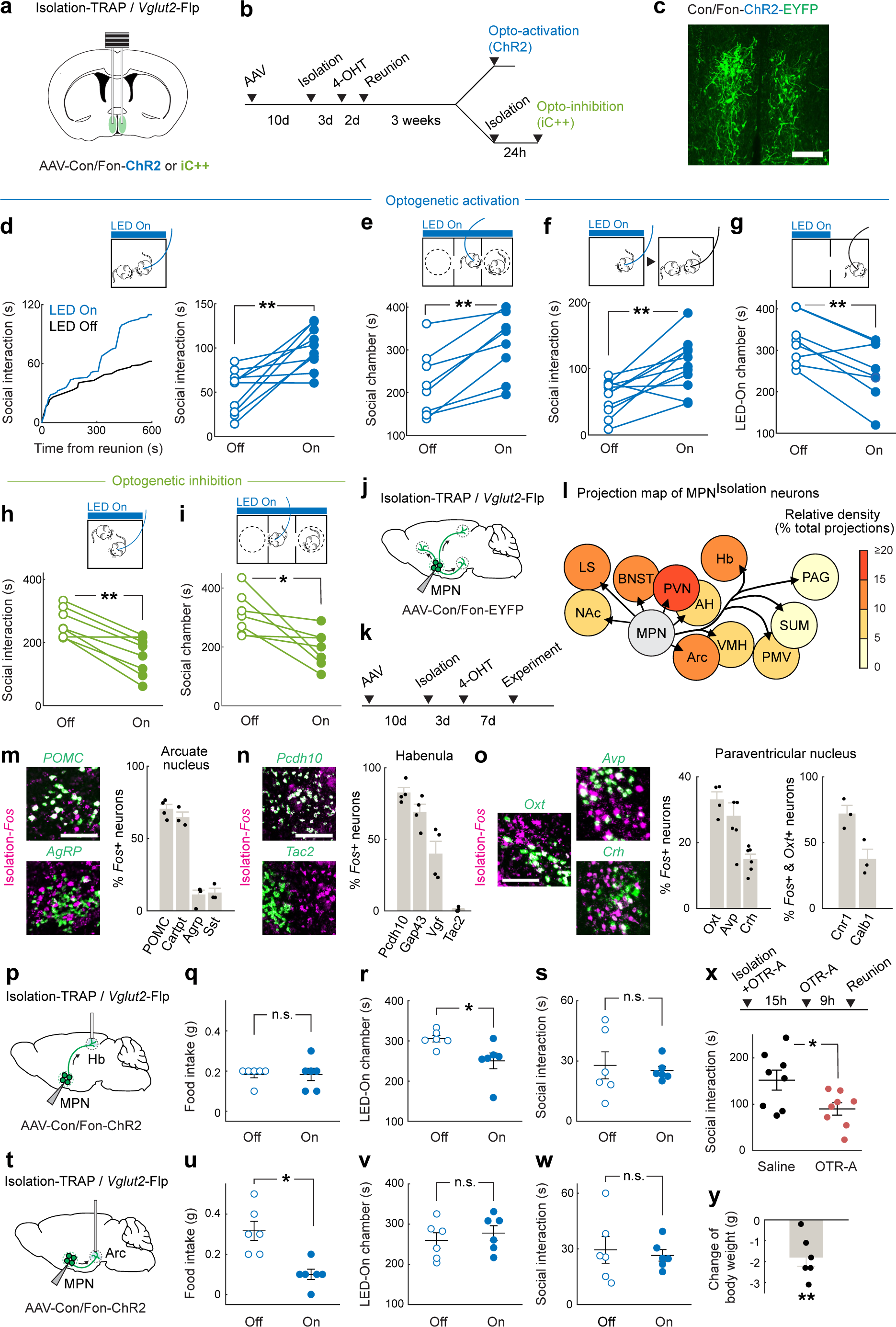
Functional characterization of MPN^Isolation^ neurons. **a**, Intersectional (Cre/Flp) strategy to target MPN^Isolation^ neurons for optogenetic manipulations. **b**, Experimental paradigms. **c**, Representative image of ChR2-expressing MPN^Isolation^ neurons. Scale bar, 200μm. **d-g**, Behavioral effects of optogenetic activation of MPN^Isolation^ neurons in different conditions: real-time stimulation in social environment, n=10 (**d**); three-chamber social preference test (**e**), n=8; pre-reunion stimulation (**f**), n=11; and real-time conditioned place preference test (**g**), n=8. **h-i**, Optogenetic inhibition of MPN^Isolation^ neurons during social reunion (**h**), n=8 and three-chamber social preference test (**i**), n=7. **j-k**, Viral tracing strategy to map projections from MPN^Isolation^ neurons. **l**, Projection map of MPN^Isolation^ neurons (NAc, nucleus accumbens. LS, lateral septum. BNST, bed nucleus of the stria terminalis. AH, anterior hypothalamus. PVN, paraventricular nucleus of hypothalamus. Hb, habenula. Arc, arcuate nucleus. VMH, ventromedial hypothalamus. PMV, ventral premammillary nucleus. PAG, periaqueductal gray. SUM, supramammillary nucleus). **m-o**, Identification of activated neurons in target regions of MPN^Isolation^ neurons. **m:** Arc, **n:** Hb and **o:** PVN. Green: marker genes, magenta: *Fos* expression following social isolation, n=3-6 for each marker gene. All scale bars, 100μm. **p-w** Behavioral effects of optogenetic activation of MPN^Isolation^->Arc and MPN^Isolation^->Hb projections on eating, real-time place preference and social behaviors, n=6. **x**, Effect of oxytocin receptor antagonist injection (i.p.) during social isolation on subsequent reunion. **y**, Changes in mouse body weight during isolation compared to group housed controls, n=6. **e**-**i, q**-**s, u**-**w, y**, Wilcoxon signed-rank tests; **x**, Mann–Whitney U test; n.s., not significant; *p<0.05, **p<0.01. All error bars represent the mean ± s.e.m.

To assess the activity of MPN^Isolation^ neurons in a cell-type specific manner, we performed microendoscopy calcium imaging of MPN*^Mc4r^*^+^ neurons in animals exposed to different behavioral regimens (Fig. 2i-q). The majority of MPN*^Mc4r^*^+^ neurons were activated by social isolation and inhibited upon social reunion (67 ± 8% of responsive neurons, ∼2.5-fold enrichment compared to pan-neuronal recording) (Fig. 2k-n and Extended Data Fig. 2d, e), and did not show significant activity in group housed animals (Fig. 2o), thus confirming their identity as the MPN^Isolation^ population. Neither tail suspension nor eating significantly modulated the activity of the MPN^Isolation^ population (Fig. 2p, q), suggesting lack of activation by general stress, salient stimuli or other physiological need. When monitoring the neural activity of MPN^Isolation^ neurons, we noticed a gradual increase of activity over minutes in the initial phase of isolation, suggesting an integration of social context (Extended Fig. 2e).

## Functional characterization of MPN^Isolation^ neurons

To examine the functional contribution of MPN^Isolation^ neurons in the emergence of social need leading to social rebound, we used optogenetic approaches to manipulate their activity and monitored corresponding behavioral changes. To specifically target MPN^Isolation^ neurons, we expressed Cre- and Flp-dependent ChR2 (Con/Fon-ChR2) in TRAP2/*Vglut2*-Flp mice in which the expression of Cre recombinase is induced by injection of 4-hydroxytamoxifen (4-OHT) during social isolation (Fig. 3a-c). Optogenetic activation of MPN^Isolation^ neurons in group housed mice whose social need is satiated induced significant social interaction (Fig. 3d), as well as enhanced social preference in a three-chamber sociability assay (Fig. 3e). Interestingly, optogenetic activation applied before the reunion step also led to a subsequent rebound in social contact (Fig. 3f), suggesting that a sustained motivational state can be triggered by transient activation of MPN^Isolation^ neurons. Place preference assays showed that mice avoided the chamber coupled with optogenetic activation (Fig. 3g), indicating that activity of MPN^Isolation^ neurons conveys negative valence, and thus may encode the aversive emotional state associated with isolation. We used a similar intersectional approach to inhibit MPN^Isolation^ neurons via expression of a light-activated chloride channel (iC++). MPN^Isolation^ neuronal inactivation significantly reduced social rebound and social preference after isolation (Fig. 3h, i). Similar results were obtained by optogenetic manipulation of MPN*^Mc4r^*^+^ neurons (Extended Data Fig. 5).

**Figure 4.**
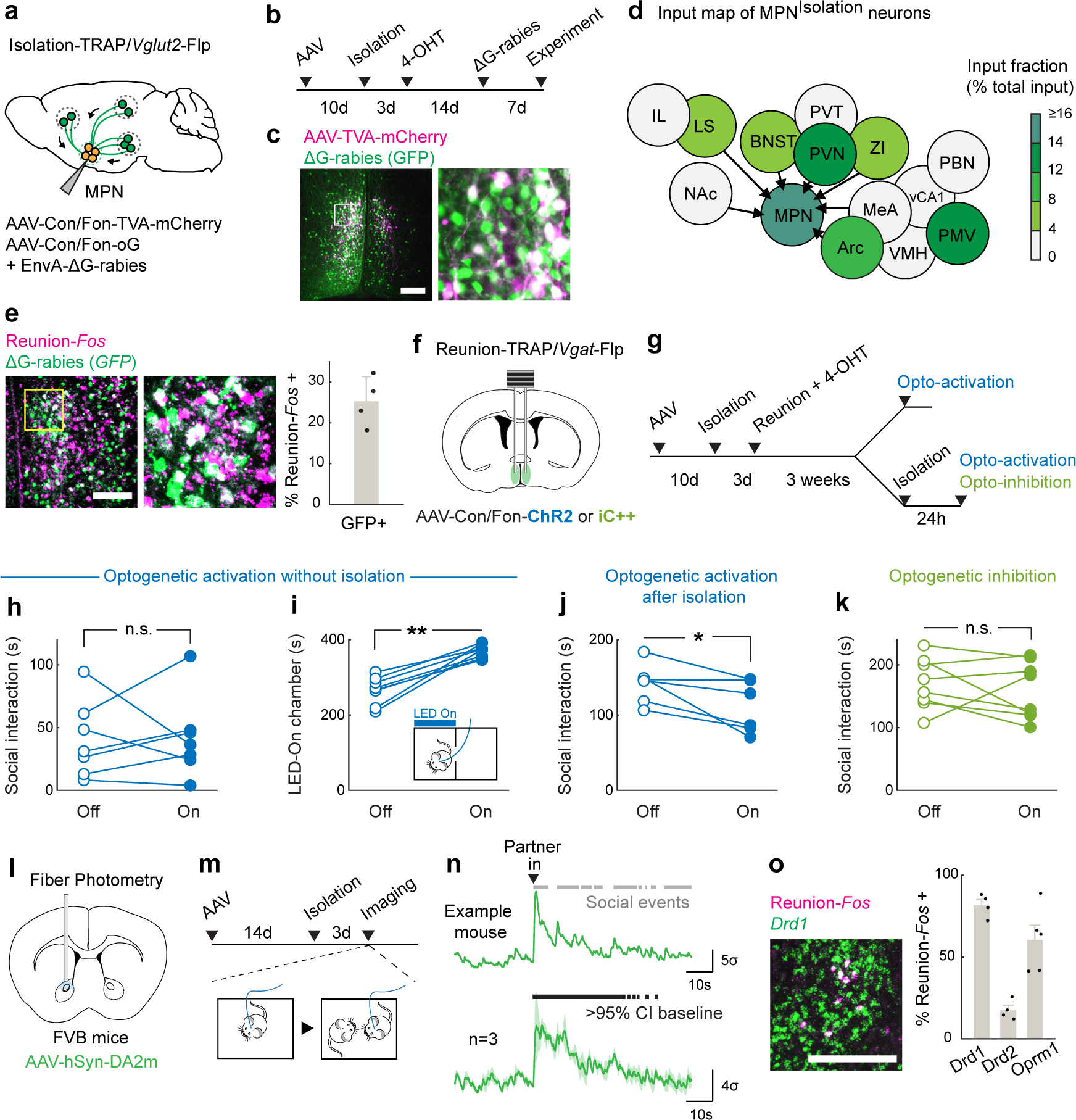
MPN^Reunion^ neurons modulate social satiety. **a-b**, Intersectional (Cre/Flp) viral tracing strategy to map input brain regions of MPN^Isolation^ neurons. **c-d**, Representative image of MPN^isolation^ starter neurons (**c**) and input brain regions (**d**). (LS, lateral septum. IL, infralimbic cortex. NAc, nucleus accumbens. BNST, bed nucleus of the stria terminalis. PVN, paraventricular nucleus of hypothalamus. ZI, zona incerta. Arc, arcuate nucleus. VMH, ventromedial hypothalamus. MeA, medial amygdala. PMV, ventral premammillary nucleus. vCA1, ventral hippocampal CA1. PBN, parabrachial nucleus) **e**, Representative image and quantification of MPN^Reunion^ neurons input into MPN^Isolation^ neurons, n=4. **f-g**, Intersectional (Cre/Flp) strategy to target MPN^Reunion^ neurons for optogenetic manipulations. **h-j**, Behavioral effects of optogenetic activation of MPN^Reunion^ neurons in different conditions: without isolation (**h**), n=7; after isolation (**j**), n=6; and real-time place preference test (**i**), n=7. **k**, Behavioral effects of optogenetic inhibition of MPN^Reunion^ neurons during social reunion, n=8. **l-m**, Experimental strategy to record dopamine release in NAc during social reunion. **n**, Dopamine release in NAc upon social reunion, n=3. Gray bars (upper) indicate social events and black bars (lower) indicate the significance of enhanced activity above 95% confidence interval (CI) of the baseline. **o**, Cell type identification of NAc neurons activated during reunion, n=4-5. **h**-**k**, Wilcoxon signed-rank tests; n.s., not significant; *p<0.05, **p<0.01. Shaded areas and all error bars represent the mean ± s.e.m. All scale bars, 200μm.

**Figure 5.**
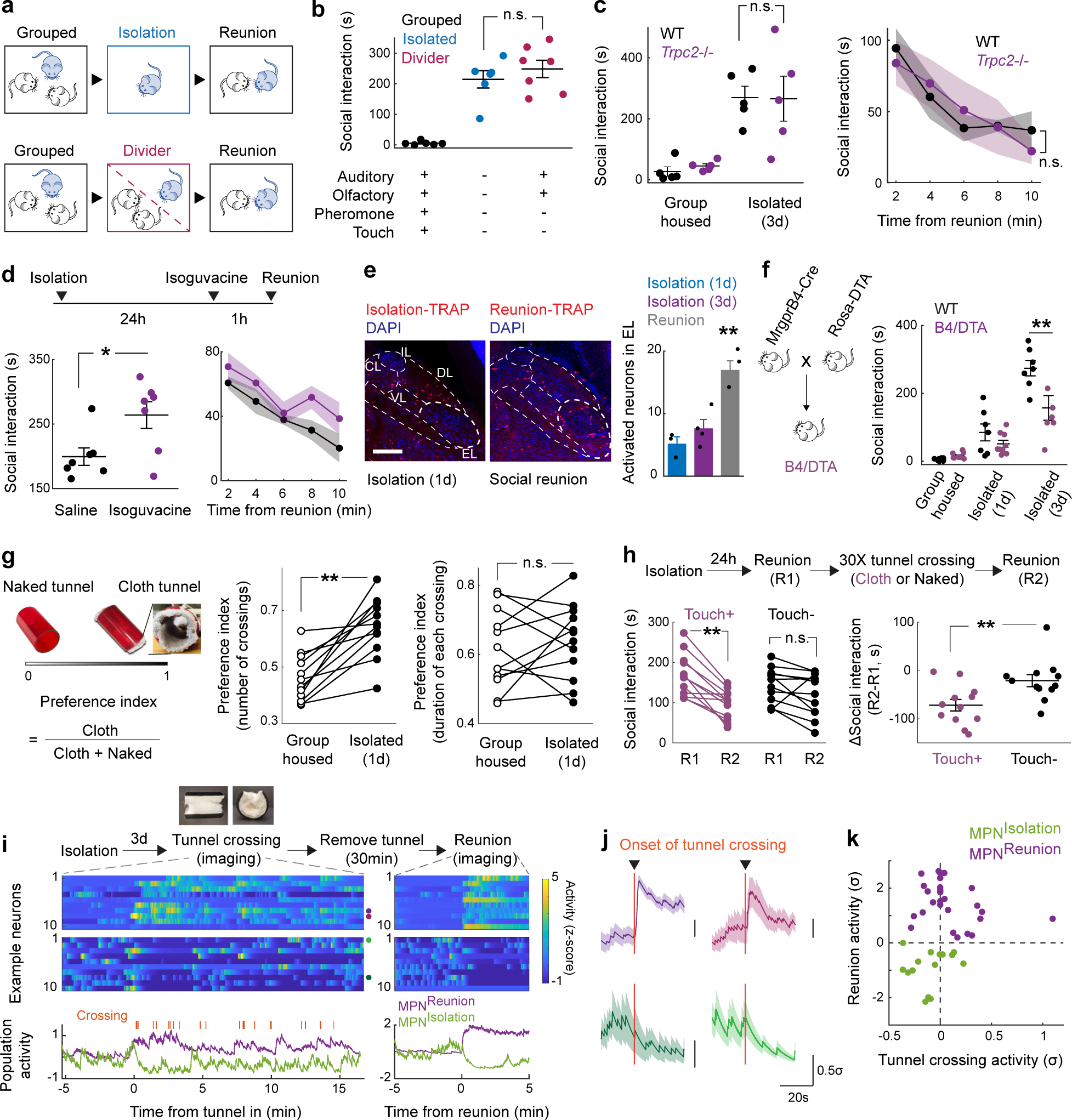
Sensory basis of social need and social satiety. **a-b**, Sensory contribution to social rebound, n=6 for standard isolation, n=7 for divider experiments, “+” and “-” indicate the presence and the absence of a given sensory modality, respectively. **c**, Social rebound and satiation in Trpc2-/- mice, n=5. **d**. Total time of social interaction following isolation with acute inhibition of touch sensation following i.p. injection of isoguvacine before social reunion, n=7. **e**, Representative images and quantification of neuronal activity in PBN subnuclei (EL, external lateral; IL, Internal lateral; CL, central lateral; VL, ventral lateral; DL, dorsal lateral) during social isolation and social reunion, n=3-4 in each condition. Scale bar, 200μm. **f**, Effect of genetic ablation of Mrgprb4 neurons using DTA on total time of social interaction following various durations of isolation, n=6-8 for each condition. **g**, Preference of mice for crossing cloth over naked tunnels during group housing or following isolation, n=12. **h**, Effect of cloth or naked tunnel crossing on social rebound, n=12. **i,** Gentle touch modulates the activity of MPN^Reunion^ (n=27) MPN^Isolation^ neurons (n=15), n=2 mice. Purple and green curves represent average activity of MPN^Isolation^ and MPN^Reunion^ neurons, respectively. **j**, Example neurons that are activated (n=2, upper) or inhibited (n=2, lower) by cloth tunnel crossing. **k,** Neuronal activity (z-scored) during social reunion and cloth tunnel crossing. Dots represent single neurons. **b, e**, Kruskal-Wallis test; **c** left, **d** left, **f, h** right, Mann–Whitney U test; **c** right, **d** right, Two-way ANOVA; **d** left, **g, h** left, Wilcoxon signed-rank tests; n.s., not significant; *p<0.05, **p<0.01. All shaded areas and error bars represent the mean ± s.e.m.

From these experiments, the activity of MPN^Isolation^ neurons appears essential to the expression of a social isolation state and is sufficient to trigger social interaction in socially satiated mice.

## MPN^Isolation^ neurons and associated neural circuits

Social isolation modulates a wide range of emotional states and behaviors^5,6^. For instance, mice find the context associated with social isolation aversive^23^ (Fig. 3g), and isolation leads to significant modulation of body weight^30^ and an increase in social seeking^3,22^. To investigate how MPN^Isolation^ neurons communicate with other brain regions to exert these functions, downstream targets were identified by viral expression of EYFP in MPN^Isolation^ neurons (Fig. 3j-l and Extended Data Fig. 6a). Projections of MPN^Isolation^ neurons were mostly found in the hypothalamus (PVN, Arc, VMH, SUM, PMV), in regions conveying aversive emotions (LS, BNST, Hb), and in downstream motor relay (PAG) (Fig. 3l and Extended Data Fig. 6a). These results were further confirmed using a conditional virus expressing synaptophysin–mRuby and membrane bound-GFP in MPN^Isolation^ neurons to delineate synapses from fibers of passage (Extended Data Fig. 6b). We then aimed to characterize in more depth the identity of activated cells in 4 of the downstream target regions, chosen based on their strong *Fos* activation during isolation and dense projections from MPN^Isolation^ neurons, namely: paraventricular nucleus of the hypothalamus, PVN; habenula, Hb; arcuate nucleus, Arc; and lateral septum, LS. Expression analysis of *Fos* after isolation together with selected marker genes enabled us to identify specific cell types in these four brain regions (Fig. 3m-o and Extended Data Fig. 6c) that may reside downstream of MPN^Isolation^ neurons and encode different aspects of the isolation state triggered by MPN^Isolation^ neuronal activity. We found that *Gap43*+, *Pcdh10*+ and *Vgf*+ neurons in the lateral habenula^31,32^, *Nts*+ and *Crhr2*+ neurons in LS^33,34^, and *Crh*+ neurons in PVN, all previously shown to process negative emotions, were activated during social isolation. We tested the function of MPN^Isolation^-to-Hb projections by expressing ChR2 in MPN^Isolation^ neurons and implanting optical fibers above Hb projections (Fig. 3p) and found that light stimulation promoted real-time avoidance (Fig. 3r) but did not affect eating or social behaviors (Fig. 3q, s). We also found that *POMC*+ and *Cartpt*+ appetite-suppressing neuronal populations in Arc were activated during social isolation (Fig. 3m), consistent with the observed body weight loss (Fig. 3y). Optogenetic activation of MPN^Isolation^ -to-Arc projections inhibited food intake (Fig. 3u) but did not induce negative valence or affect social interaction (Fig. 3v, w), suggesting a direct inhibition of eating drive by social need. The distinct behavioral effects elicited by stimulation of MPN^Isolation^ -to-Hb and MPN^Isolation^ to-Arc projections rule out possible cross-talks between MPN^Isolation^ projections via backpropagated action potentials induced by terminal activation. We also found that social isolation activated *Oxt*+, *Avp*+ neurons in PVN and *Oxtr*+ neurons in LS (Fig. 3o and Extended Data Fig. 6c) which may underlie the enhanced social drive triggered by isolation. Indeed, blocking oxytocin receptor (OTR) during isolation by intraperitoneal injections of OTR antagonist (OTR-A) significantly reduced social rebound (Fig. 3x).

**Figure 6.**
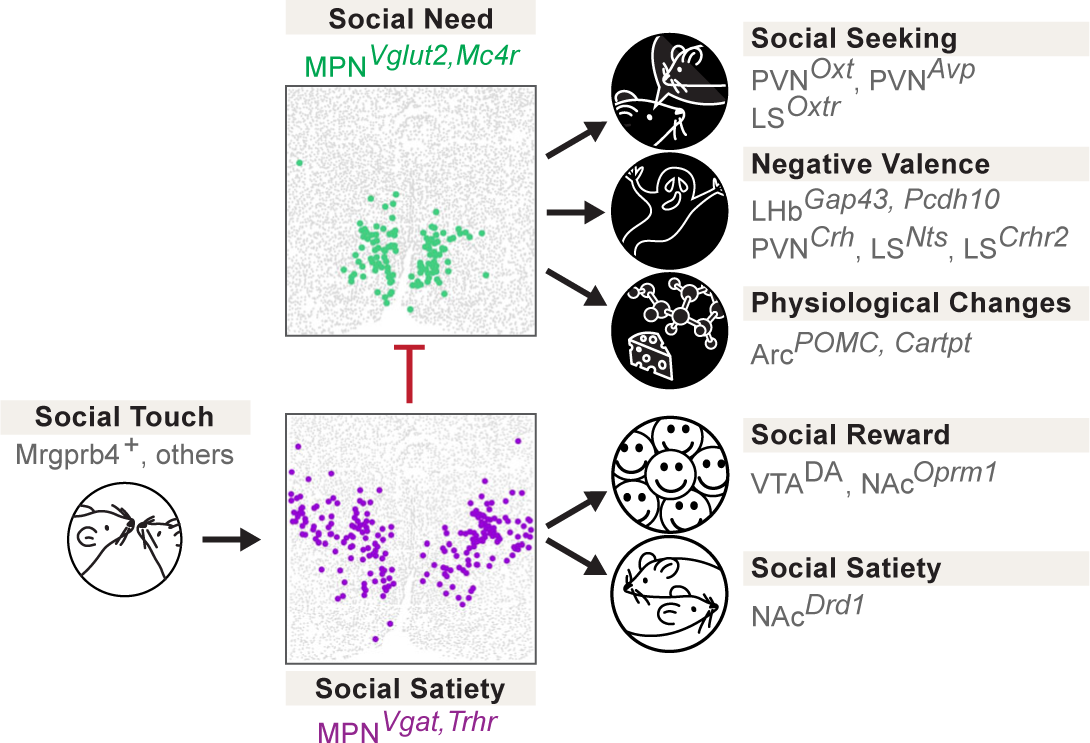
Model: neural circuits underlying dynamic control of social need. Gentle touch associated with social interaction leads to the activation of MPN^Reunion^ neurons, the recruitment of social reward circuits, inhibition of MPN^Isolation^ neurons and social satiety. Conversely, absence of social touch input during isolation inactivates MPN^Reunion^ neurons and in turn dis-inhibits (activates) MPN^Isolation^ neurons, which enhances social motivation, negative valence and other physiological functions that constitute the isolation state.

## MPN^Reunion^ neurons and the modulation of social satiety

MPN^Isolation^ neurons showed a quick suppression of activity upon reunion (Fig. 2c, d, k-m), thus signaling the end of social isolation. To better understand the circuit mechanisms underlying the observed inhibition of MPN^Isolation^ neurons upon social contact, we performed monosynaptic retrograde tracing^35^ from MPN^Isolation^ neurons. The TVA receptor and optimized G protein were expressed in MPN^Isolation^ neurons using the intersectional strategy described earlier (Fig. 4a). Two weeks later, G-deleted rabies virus was injected into the MPN and brain-wide retrograde rabies signals were measured (Fig. 4a-d). This approach identified 12 input areas for MPN^Isolation^ neurons (Fig. 4d and Extended Data Fig. 6d), many overlapping with MPN^Isolation^ projection areas, thus revealing extensive recurrent connectivity within the isolation circuit. Among MPN^Isolation^ input areas we noticed a strong local signal within the preoptic area. This observation prompted us to examine the identity of local input neurons and assess whether MPN^Reunion^ neurons are a direct source of inhibition of MPN^Isolation^ neurons upon reunion. We examined the co-expression of presynaptic rabies (GFP) from MPN^Isolation^ neurons and *Fos* induced by social reunion, and observed a significant overlap (Fig. 4e), suggesting that MPN^Reunion^ neurons directly synapse onto MPN^Isolation^ neurons. Since ∼80% of MPN^Reunion^ neurons are GABAergic, their activation likely directly suppresses the activity of MPN^Isolation^ neurons during social rebound, thus serving as a social satiety signal.

Next, we targeted MPN^Reunion^ neurons by expressing Con/Fon-ChR2 in TRAP2/*Vgat*-Flp mice and inducing the expression of Cre recombinase by injection of 4-hydroxytamoxifen (4-OHT) during social reunion (Fig. 4f, g). Optogenetic activation of MPN^Reunion^ neurons did not affect social interaction in group housed animals (Fig. 4h). However, activation of these neurons during social reunion after a period of isolation led to a decreased social rebound (Fig. 4j). These results support the hypothesis that activity of MPN^Reunion^ neurons provides a social satiety signal when isolated animals are reunited. Importantly, in a real-time place preference assay, mice preferred to stay in the chamber associated with optical stimulation implying a positive valence conveyed by the activity of MPN^Reunion^ neurons (Fig. 4i). Inhibition of MPN^Reunion^ neurons during reunion did not further increase the rebound, perhaps due to the incomplete capture of MPN^Reunion^ neurons by the TRAP approach or the existence of other social satiation mechanisms (Fig. 4k).

To gain a broader understanding of the neural circuitry underlying social homeostasis, we further mapped the input and output of MPN^Reunion^ neurons and identified brain regions, such as LS, NAc, BNST, PVN and Arc that overlap with circuits associated with MPN^Isolation^ neurons (Extended Data Fig. 7), suggesting a shared neural network in the regulation of social need and social satiety. MPN^Reunion^ neurons showed dense projections to the ventral tegmental area (VTA), which has robust dopaminergic projections to NAc shown to be essential in the control of social reward^21^. We virally expressed the dopamine sensor GRAB_DA2m_^36^ in NAc, measured fluorescence changes with fiber photometry, and observed significant dopamine release in NAc upon social reunion (Fig. 4l-n). Since both MPN^Isolation^ and MPN^Reunion^ neurons receive inputs from NAc, we speculate that MPN-VTA-NAc circuit may provide real-time modulation of social motivation during social reunion. Using *in situ* hybridization, we further identified reunion-activated NAc neurons as expressing the dopamine D1 receptor (D1R) as well as the opioid receptor *Oprm1* (Fig. 4o), shown to mediate social reward^37^. NAc D1R neurons are inhibitory and may provide negative feedback^38^ to both MPN^Isolation^ and MPN^Reunion^ neurons, thus contributing to the decay of social rebound during reunion.

These data identify a brain-wide neural circuitry underlying social need regulation. Social stimuli activate MPN^Reunion^ neurons, which further recruit downstream social reward circuits and inhibit MPN^Isolation^ neurons. Conversely, during social isolation, absence of sensory input silences MPN^Reunion^ neurons and in turn dis-inhibits MPN^Isolation^ neurons, which further triggers negative valence, social motivation and other physiological changes associated with social isolation.

## Sensory basis of social need and social satiety

How do animals assess their social environment and discern if they are alone or together? To explore this issue, we investigated the contribution of various senses to the emergence and satiation of social need. In an initial experiment, mice were separated from siblings in their home cage by a perforated divider such that auditory and olfactory cues could be sensed by animals across the divider (Fig. 5a). Since FVB mice become blind by weaning age, the role of visual cues was not addressed. Surprisingly, a significant social rebound, comparable to the rebound induced in single housed mice, was observed in the reunion period following 3 days of separation (Fig. 5b). This suggests that signals crossing the divider — auditory, and olfactory cues — are not sufficient to fulfill the social need of the separated mice. We therefore tested two of the remaining sensory modalities that require direct contact, namely pheromone sensing and touch, as candidate cues enabling mice to perceive social context. The potential contribution of pheromone sensing was assessed by measuring the behavior of *Trpc2*-/- mice, which are genetically impaired in vomeronasal pheromone sensing^39^. *Trpc2*-/- mice displayed normal social rebound after isolation and satiation during reunion (Fig. 5c), suggesting that, instead, somatosensation may represent the essential sensory modality for mice to assess the presence or absence of others, and thus either lead to social satiety or social need, respectively.

To assess the contribution of touch to social need and social satiation more directly, we first examined the effect of acute tactile inhibition by intraperitoneal injection of isoguvacine, a peripherally restricted GABA_A_ receptor agonist that attenuates peripheral mechanosensory neuron firing^40^, just before social reunion. Mice treated with isoguvacine showed prolonged social rebound (Fig. 5d), suggesting that attenuated mechanosensory neuron signaling may delay the satiation of social need. In parallel, we examined the neuronal activity induced by social rebound in the lateral parabrachial nucleus (PBN_L_), a brain region densely innervated by the axons from spinoparabrachial (SPB) neurons that convey touch information from the spinal cord to the brain^41^. Interestingly, we found that social rebound robustly activates the external lateral subnucleus of PBN (PBN_EL_) (Fig. 5e and Extended Data Fig. 8a) which was previously implicated in affective touch^41^. PBN_EL_-activating SPB neurons receive direct synaptic input from *Mrgprb4*+ mechanosensory neurons^41^, and *Mrgprb4*+ neurons are implicated in mediating pleasant and social touch in mice^42,43^, prompting us to examine whether lack of *Mrgprb4*+ neurons influences social rebound. After crossing *Mrgprb4*-Cre mice to a Cre-dependent diphtheria toxin subunit A (DTA) line, we found that mice with ablated *Mrgprb4*+ neurons (B4/DTA) showed a slight but significant reduction in social rebound compared to wildtype controls (Fig. 5f). This result suggests that constitutive loss of at least one neuronal population previously implicated in affective touch leads to reduced sensitivity to social environment and thus hampers the generation of social drive during isolation. The reduced but significant rebound still observed in B4/DTA mice (Fig. 5f) suggests that other types of somatosensory fibers are also involved in the sensation of social context.

Inspired by Harlow’s pioneering work^44^, in which separated infant rhesus monkeys strongly preferred and attached to a soft cloth mother rather than a rigid wire surrogate, we designed a mouse version of a comfort-touch preference assay. In this assay, mice were exposed to two types of plastic tunnels, one internally coated with soft cloth material (cloth tunnel) and the other left naked (naked tunnel) to serve as a control (Supplementary Video 3). After isolation, but not when group housed, FVB mice showed a significant preference for crossing the cloth versus naked tunnel as quantified by the relative number of crossings of each tunnel (Fig. 5g, left graph). The relative duration of crossing the cloth vs naked tunnel remained unchanged between group housing and isolation conditions (Fig. 5g, right graph), suggesting that the preference of the cloth tunnel results from dynamic gentle touch stimulation by the cloth lining rather than shelter or warmth seeking. Interestingly, strains that show low rebound behaviors, such as C57 mice, did not display such touch preference (Extended Data Fig. 8b). To further investigate whether comfort touch could at least partially fulfill social need, we let isolated mice cross through either cloth or naked tunnels 30 times and measured the rebound intensity before and after tunnel crossing. Remarkably, soft-touch-stimulated mice displayed significantly lower rebound after tunnel crossing, while mice running through naked tunnels had similar rebound as prior to the tunnel crossing (Fig. 5h). Similarly, co-housing with the cloth tunnel, but not with the naked tunnel, during isolation reduced social rebound (Extended Data Fig 8c).

How does soft touch affect hypothalamic circuits underlying social need and social satiety and does it directly modulate the activity of MPN^Reunion^ and MPN^Isolation^ neurons? To answer these questions, we modified the cloth tunnel by adding an opening on top such that mice can run through such tunnel with a head implant and attached wire during microendoscopy calcium imaging. After three days of isolation, the implanted mice were allowed to freely access the cloth tunnel for 15 minutes (10-20 crossings) while the MPN neuronal activity was recorded. Then the tunnel was removed for 30 minutes, and a 5-minute reunion assay was applied, enabling us to identify MPN^Reunion^ or MPN^Isolation^ neurons based on their activation or inhibition during social reunion. We then examined the activity of these two populations during the previous occurrences of cloth tunnel crossing. Strikingly, over 90% of MPN^Isolation^ neurons were inhibited and ∼35% of MPN^Reunion^ neurons were excited during cloth tunnel crossing (Fig. 5i-k). These results indicate that gentle touch is effective in quenching MPN^Isolation^ and eliciting MPN^Reunion^ neuronal activity, thus mimicking signals underlying social satiation. Overall, these results suggest that physical touch is a key sensory channel for the perception of social environment such that a lack of touch sensation leads to the emergence of social need, while its presence provides social satiety.

## Discussion

In this study, we have uncovered two molecularly defined and interconnected neuronal populations in the medial preoptic nucleus of the hypothalamus that form a key regulatory node underlying social homeostasis (Fig. 6). Excitatory MPN^Isolation^ neurons marked by *Slc17a6* (*Vglut2)*, *Mc4r* and *Cartpt*, are active during social isolation and inhibited upon social reunion. Their activity drives social interaction and is associated with negative valence, while their silencing inhibits social rebound that follows a period of social isolation, suggesting a key role in encoding social need when animals are socially deprived. By contrast, inhibitory MPN^Reunion^ neurons marked by *Slc32a1* (*Vgat)* and *Trhr* are activated during social reunion and inhibited by social isolation. Their activity reduces social interaction and is associated with positive valence, suggesting a role in encoding social satiety when animals are reunited after isolation. Activity imaging and connectivity analysis showed that MPN^Reunion^ neurons are activated by gentle touch stimuli and send direct inhibitory synaptic input to MPN^Isolation^ neurons. These findings suggest a circuit mechanism in which loss of social contact during a period of isolation triggers the activity of MPN^Isolation^ neurons to signal an aversive state and enhance social motivation through MPN^Isolation^ projections to LHb*^Gap43^*, PVN^Crh^, LS^Crhr2^ and PVN*^Oxt^* neurons. By contrast, social contact after isolation drives social reward and satiety through MPN^Reunion^ neurons via the VTA-NAc circuit, leading to an increase in DA release.

Strikingly, the opposite functions and mutual interaction demonstrated between MPN^Reunion^ and MPN^Isolation^ neurons bare strong similarities with the organization of known hypothalamic circuits underlying the homeostatic control of physiological needs, such as the opposite function of Arc*^Agrp^* and Arc*^POMC^* neurons in appetite control^13,14^, and similar organization for thirst^15-17^ and sleep^18^. Intriguingly, unlike that of food and water, social need is typically not considered as a survival need. Thus, the similar neural architectures identified in the control of social and physiological needs may highlight the significance of social drive for survival as well as reflect a common strategy for evolutionarily conserved behavioral drives.

Physiological needs are monitored through the detection of peripheral signals, such as metabolite concentration for appetite^13,14^ and osmolality for thirst^15-17^. Our data suggest that touch is a key signal conveying the social environment (Fig. 5). This observation is in line with recent studies showing that gentle tactile stimulation in rodents significantly modulates the motivation of social behaviors^45-47^, and that zebrafish sense the presence of conspecifics via mechanoreceptors in the lateral line organ^48^. In our study, inhibition of touch sensation during social reunion impaired the satiation of social drive and led to prolonged social rebounds. Constitutive ablation of sensory neurons expressing *Mrgprb4,* a population shown to contribute to social touch^42,43^, led to reduced social rebound. The effect of ablating *Mrgprb4* neurons was significant but relatively modest, thus implicating other peripheral mechanosensory neuron populations in social touch. Social isolation effectively promotes touch seeking behaviors, which in turn inhibits MPN^Isolation^ neurons and reduces social need. Social touch has been shown to play an essential role in brain development, stress alleviation, attention improvement and pain relief, while social touch avoidance is a hallmark of neurodevelopmental disorders such as autism spectrum disorder (ASD)^40^. Recent findings have identified sensory neuron types and associated neural circuits mediating social touch^42,43,49^. Our work further emphasizes how social touch is an essential contributor to social homeostasis.

Social touch is rewarding and triggers dopamine release in the nucleus accumbens (NAc)^43^, thus contributing to the rewarding nature of social interactions^19-21^. Similarly, we observed significant dopamine release in NAc upon social reunion. We also found that activation of MPN^Reunion^ neurons induces conditioned place preference and that MPN^Reunion^ neurons have extensive connectivity with VTA, which may directly trigger dopamine release during social interaction. Moreover, we observed that NAc neurons are activated during social reunion and both MPN^Isolation^ and MPN^Reunion^ neurons receive inputs from NAc, leading us to hypothesize that the MPN-VTA-NAc circuit may form a negative feedback loop that modulates MPN activity and mediates the decay of social rebound upon reunion. By contrast, social isolation activates brain areas and cell types signaling negative valence (Fig. 6), together generating an aversive emotional state. In a recent study, dopamine neurons within the dorsal raphe nucleus (DRN) were proposed to represent an aversive state associated with social isolation^22^, or alternatively arousal to salient stimuli^50^. Our tracing results indicate that MPN^Reunion^ neurons project to the DRN (Extended Data Fig. 7d, e) suggesting possible modulation of DRN by MPN^Reunion^ neurons during social reunion.

How does the brain track the duration of social isolation and trigger a scalable social rebound after increasing lengths of isolation? The duration of social isolation may be directly encoded by changes in the firing rate or population activity patterns of MPN^Isolation^ neurons. Alternatively, the activity of MPN^Isolation^ neurons may be further integrated in downstream effectors across days of isolation, and, for example, may lead to the release of neuropeptides or other signaling molecules whose concentration accumulates during isolation. It is also possible that the activity of MPN^Isolation^ neurons facilitates synaptic plasticity of local or global neural circuits, that magnify social stimuli and trigger social rebound during reunion. These hypotheses can be tested in future experiments through long-term recordings that can track ensembles of neurons across days.

Altogether, we have uncovered a brain-wide circuit centered around two hypothalamic nodes, MPN^Isolation^ and MPN^Reunion^ neurons, that monitor touch signals to provide animals with a dynamic neural representation of the social environment and underlie the behavioral and emotional control of social homeostasis. These insights into the nature and identity of circuits controlling social need and the remarkable similarity with other physiological homeostatic controls open new avenues for the understanding and treatment of mental and physical disorders induced or exacerbated by social isolation.

## Acknowledgements

We thank S. Sullivan for assistance with mouse husbandry and genotyping; S. Qian, J. Garzon, T. Han, N. H. Kaplan, Xia, I. Sabbarini and Z. Sullivan for help with experiments and analysis; R. Hellmiss and J. Mancini at MCB Graphics for help with illustrations; Dr. Connie Cepko for providing *Pde6b^rd1^* mice in C57BL/6J background; Inscopix for technical support; members of the Dulac and the Uchida laboratories for helpful advice on experiments and analysis and comments on the manuscript. This work was supported by funding from the NOMIS Foundation, Hock E. Tan and K. Lisa Yang Center for Autism Research at Harvard University, Jane Coffin Childs Medical Research Award 61-1665 to D.L., Lee and Ezpeleta funded undergraduate fellowship to A.J., and Simons Foundation Award 673021 and NIH award R01NS116593 to C.D. C.D. and D.D.G. are investigators at the Howard Hughes Medical Institute.

## Author contributions

D.L. and C.D. conceived and designed the study. D.L. performed social behavior screening/characterization, cell-type identification, optogenetics and neural tracing experiments. D.L. and M.R. performed microendoscopic calcium imaging and analysis. D.L. and A.J. performed sensory and touch related experiments. D.L and I.T.-K. performed fiber photometry and analysis. N.P. helped with optogenetic experiments. M.T developed the behavior recording system. B.L.L. analyzed scRNAseq and MERFISH data for cell type identification. S.F. assisted with in situ hybridization experiments and analysis. D.L. and S.C. assessed neuronal activity in PBN. A.C. helped with USV analysis. M.U. and N.U. provided fiber photometry setup. I.A.-S. provided Mrgprb4/DTA mouse line and shared unpublished information. C.D., M.W.-U., N.U. and D.D.G. provided instruction and comments during the research. D.L. and C.D. wrote the manuscript with input from all authors.

## Competing interests

The authors declare no competing interests.

---

**Extended Data Figure 1.**
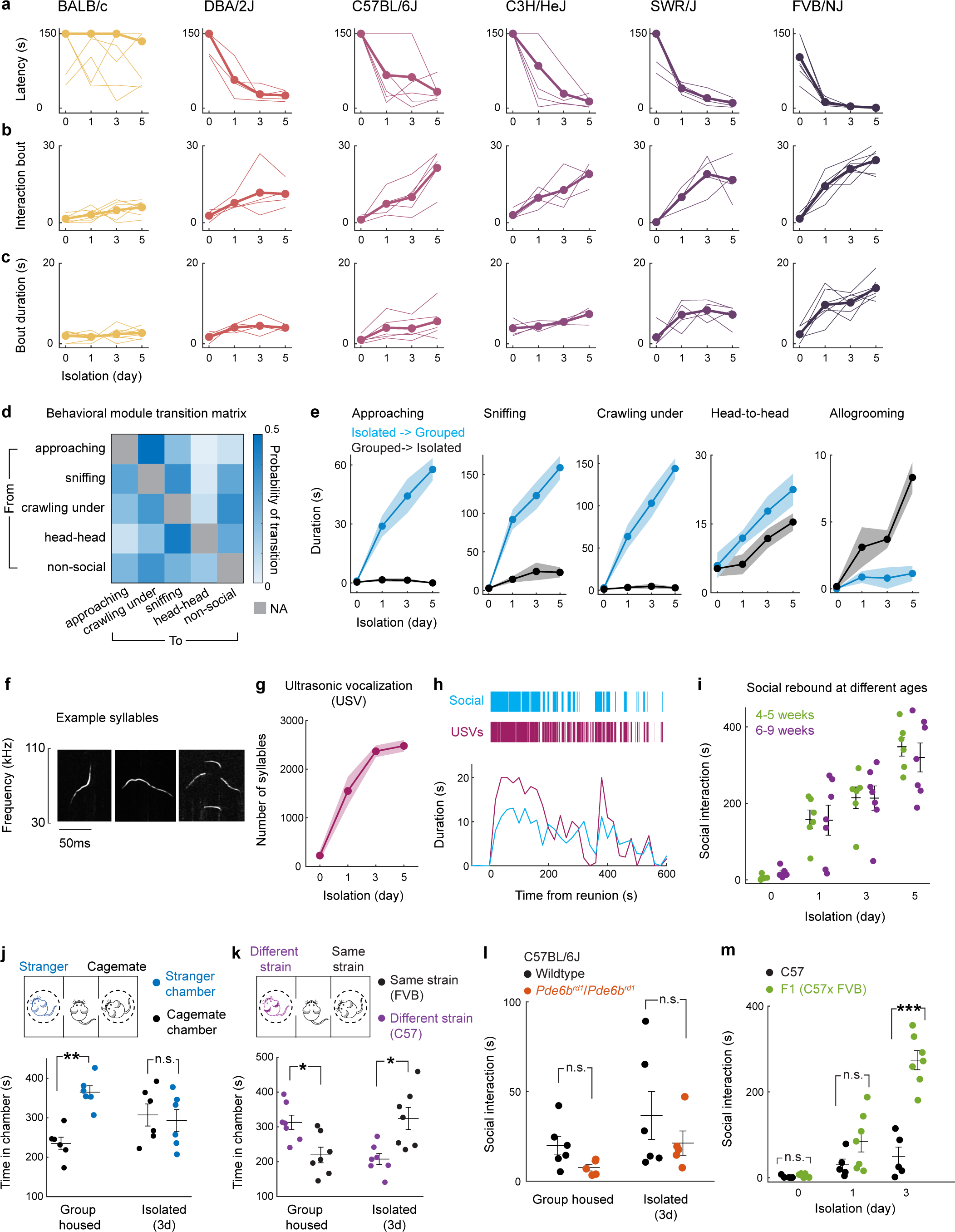
Behavioral characterization of social rebound. **a-c**, Social interaction latency, bout number and mean bout duration during social reunion. Thin lines represent individual mice, thick lines are cohort average (BALB/c n=7, DBA/2J n=4, C57BL/6J n=5, C3H/HeJ n=4, SWR/J n=4, FVB/NJ n=7). **d**, Probability matrix of transitions between behavioral modules during social rebound. **e**, Display of specific behavioral modules in paired isolated with group housed (blue) vs group housed with isolated (black) FVB mice, n=7. **f**, Example ultrasonic vocalization (USV) syllables of FVB mice recorded during social reunion. **g**, Number of USV syllables during social reunion in FVB mice after various lengths of isolation, n=4. **h**, Time course of social events and USVs from example FVB mouse. **i**, Social rebound in FVB mice of different ages, 4–5 weeks, n=6; 6-9 weeks, n=7. **j**, Three-chamber preference test between cage mate and stranger, in group housed vs isolated FVB mice, n=6. **k**, Preference test between mice from same (FVB) or different (C57) strain, n=7. **l**. Social interaction in group housed and isolated wildtype (n=6) vs *Pde6b^rd1^/Pde6b^rd1^*(n=5) C57BL/6J mice. **m**, Social interactions in C57BL/6J (C57, n=5) and in offspring of FVB/NJ x C57BL/6J mice (F1, n=7). **f**, Mann–Whitney U test; **h**, Friedman test; **j, k**, Wilcoxon signed-rank tests; **l, m**, Mann–Whitney U test; n.s., not significant; *p<0.05, **p<0.01, ***p<0.001. All shaded areas and error bars represent the mean ± s.e.m.

**Extended Data Figure 2.**
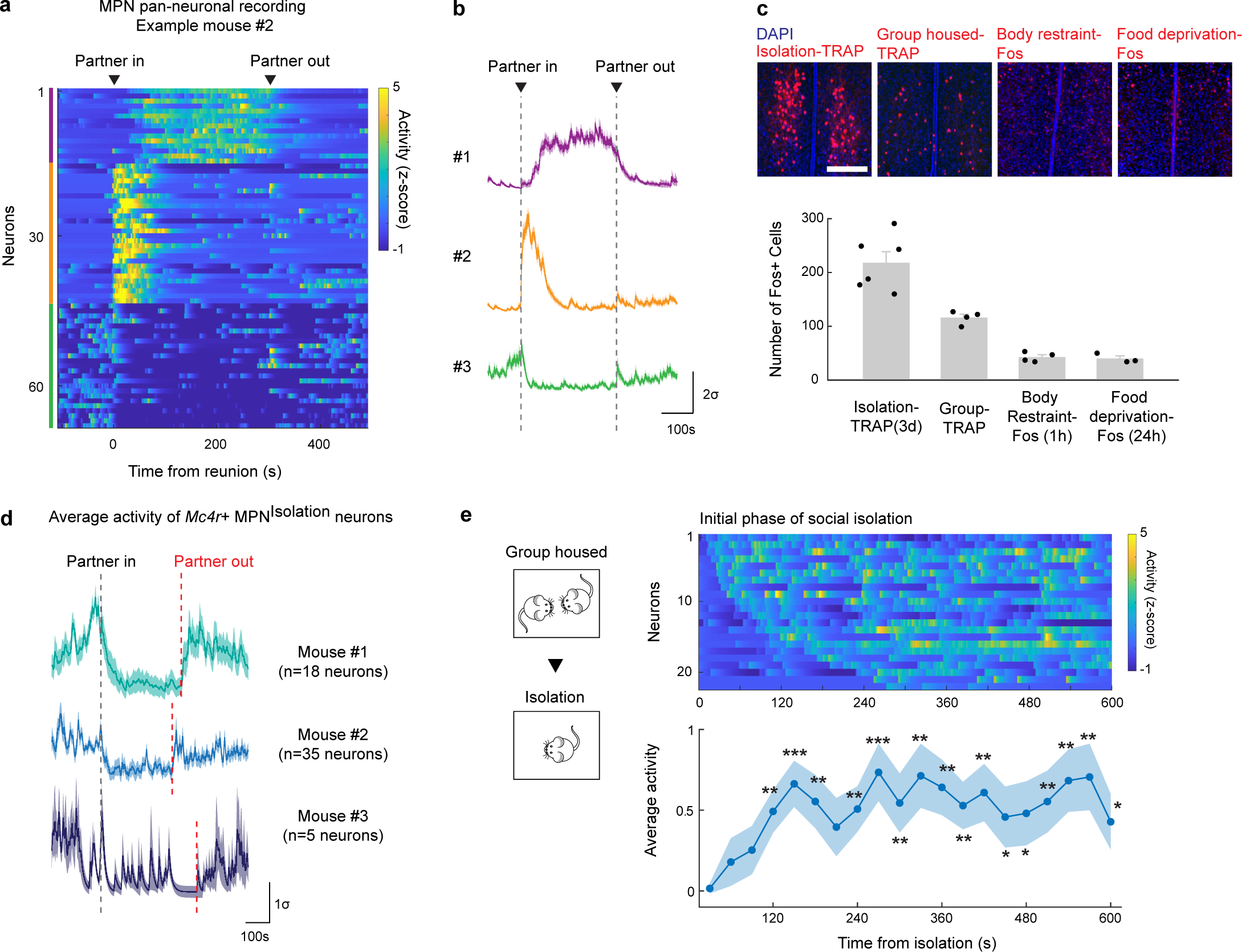
Microendoscopy Calcium imaging in MPN. **a,b** Second example mouse with MPN pan-neuronal calcium imaging during social isolation and social reunion, and 3 clusters of neurons displaying distinct activity patterns during reunion. **c**. Number of activated MPN neurons during social isolation, group housing, restraint stress (1h) and food deprivation (24h), n=3-6. Scale bar, 200μm. **d**. Average calcium activity of MPN*^Mc4r^*^+^ neurons that were inhibited during reunion, n=3. **e**. Calcium imaging of MPN^Isolation^ neurons in the initial phase of social isolation. Data below represent average activity across 27 neurons pooled from 2 mice. Wilcoxon signed-rank tests were used to estimate statistical significance of enhanced neuronal activity in each time bin (30s) vs the first time bin, *p<0.05, **p<0.01, ***p<0.001.

**Extended Data Figure 3.**
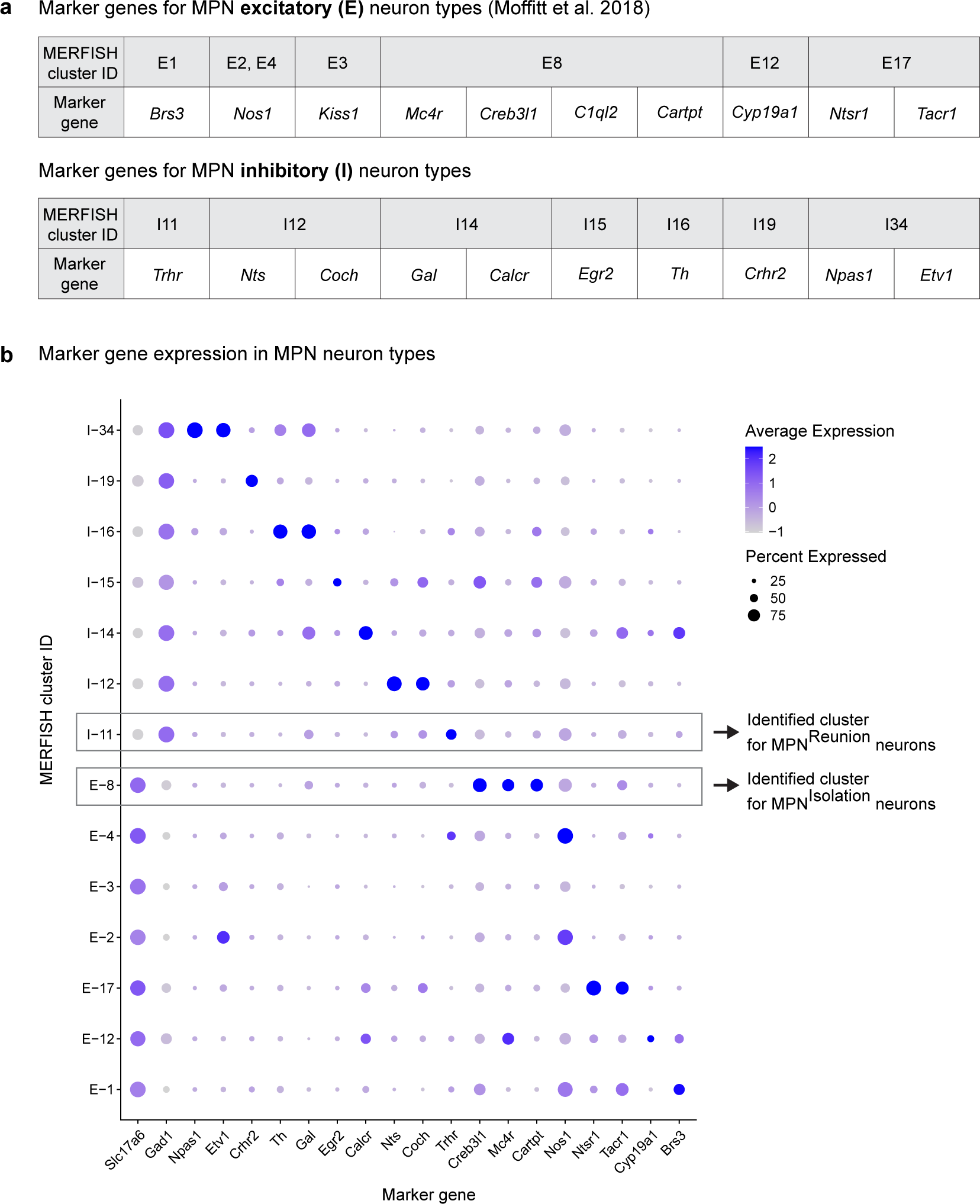
Marker genes of MPN cell types. **A**, Marker genes for different MPN excitatory and inhibitory neuronal populations based on Moffitt et al. 2018. **b**, Expression patterns of marker genes in different MPN clusters based on MERFISH experiments^29^. Boxes identify clusters that are best matched with MPN^Isolation^ and MPN^Reunion^ neurons.

**Extended Data Figure 4.**
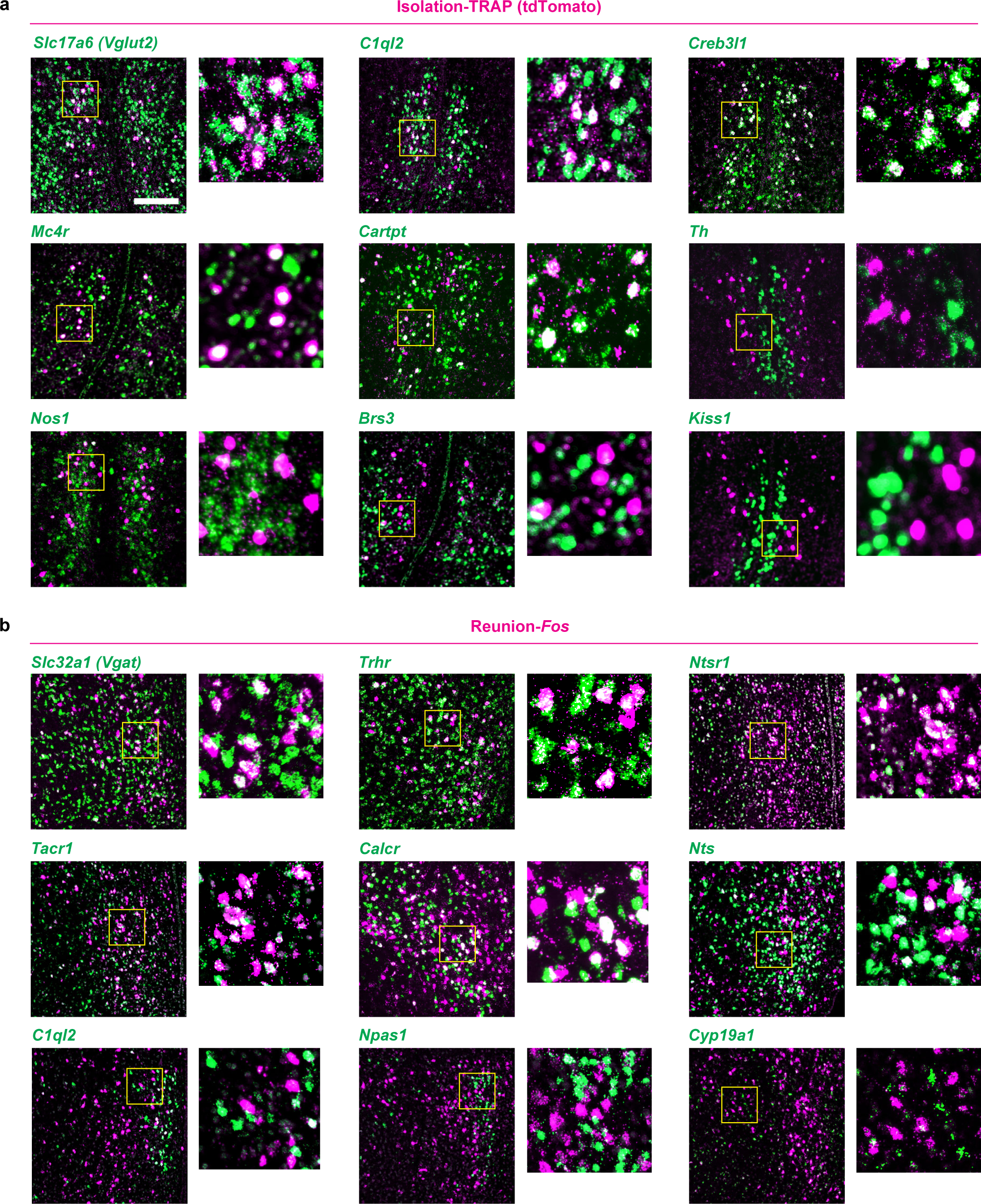
Cell type identification of MPN^Isolation^ and MPN^Reunion^ neurons. **a, b** Representative images of marker gene expression of distinct MPN cell-type cluster and overlap with MPN^Isolation^ (**a**) and MPN^Reunion^ (**b**) neurons using *in situ* hybridization. Green: marker genes for specific neuron types, magenta: *Fos* expression during social isolation (**a**) or social reunion (**b**). Scale bar, 200μm (all images). Zoom-in box, 150μm x 150μm.

**Extended Data Figure 5.**
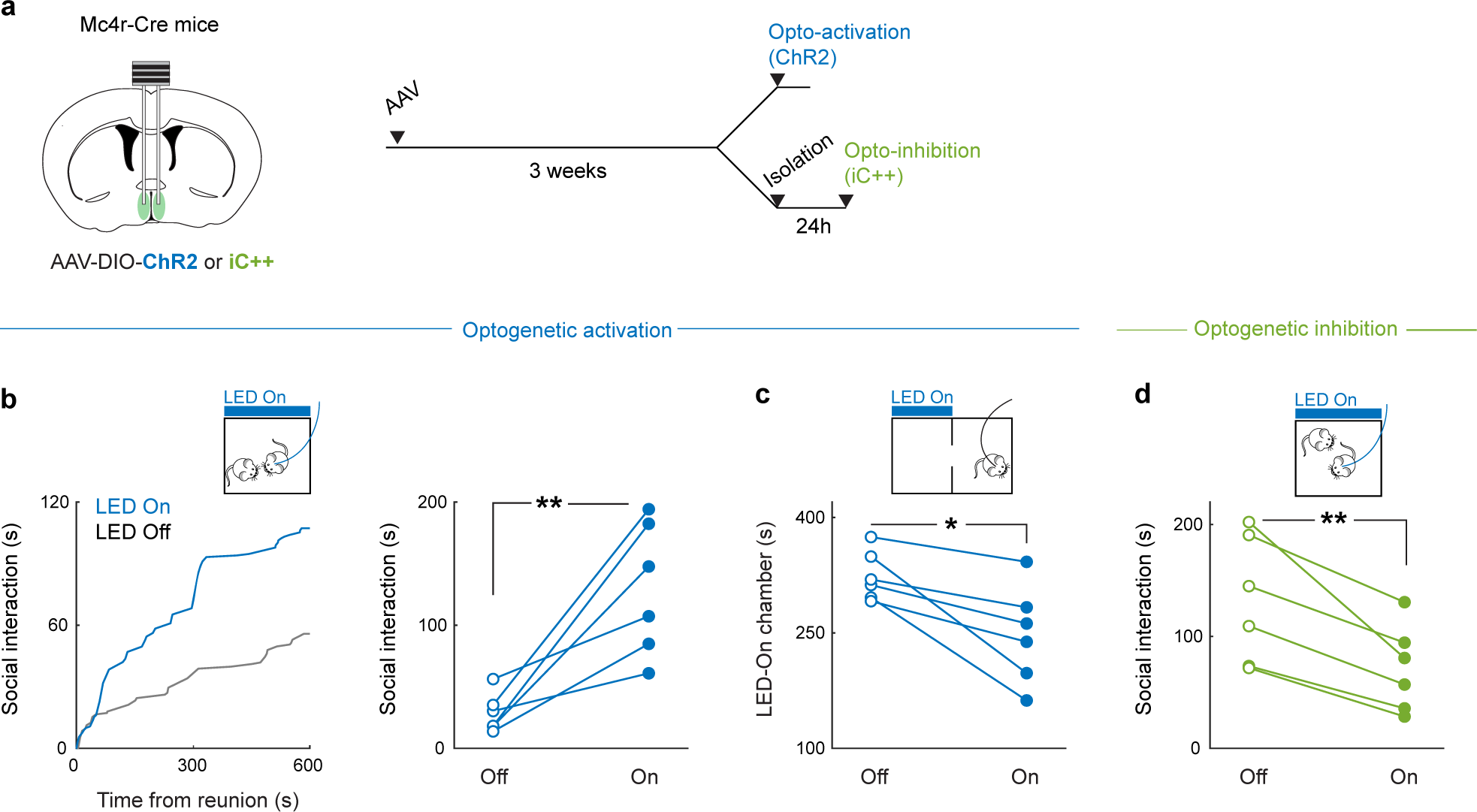
Optogenetic manipulation of MPNMc4r^+^ neurons. **a**, Strategy to target MPN*^Mc4r^*^+^ neurons for optogenetic manipulations. **b-c,** Behavioral effects of optogenetic activation of MPN*^Mc4r^*^+^ neurons during social interaction and real-time place preference test, n=6. **d**. Behavioral effects of optogenetic inhibition of MPN^Mc4r+^ neurons during social reunion, n=6. **b**-**d**, Wilcoxon signed-rank tests; *p<0.05, **p<0.01.

**Extended Data Figure 6.**
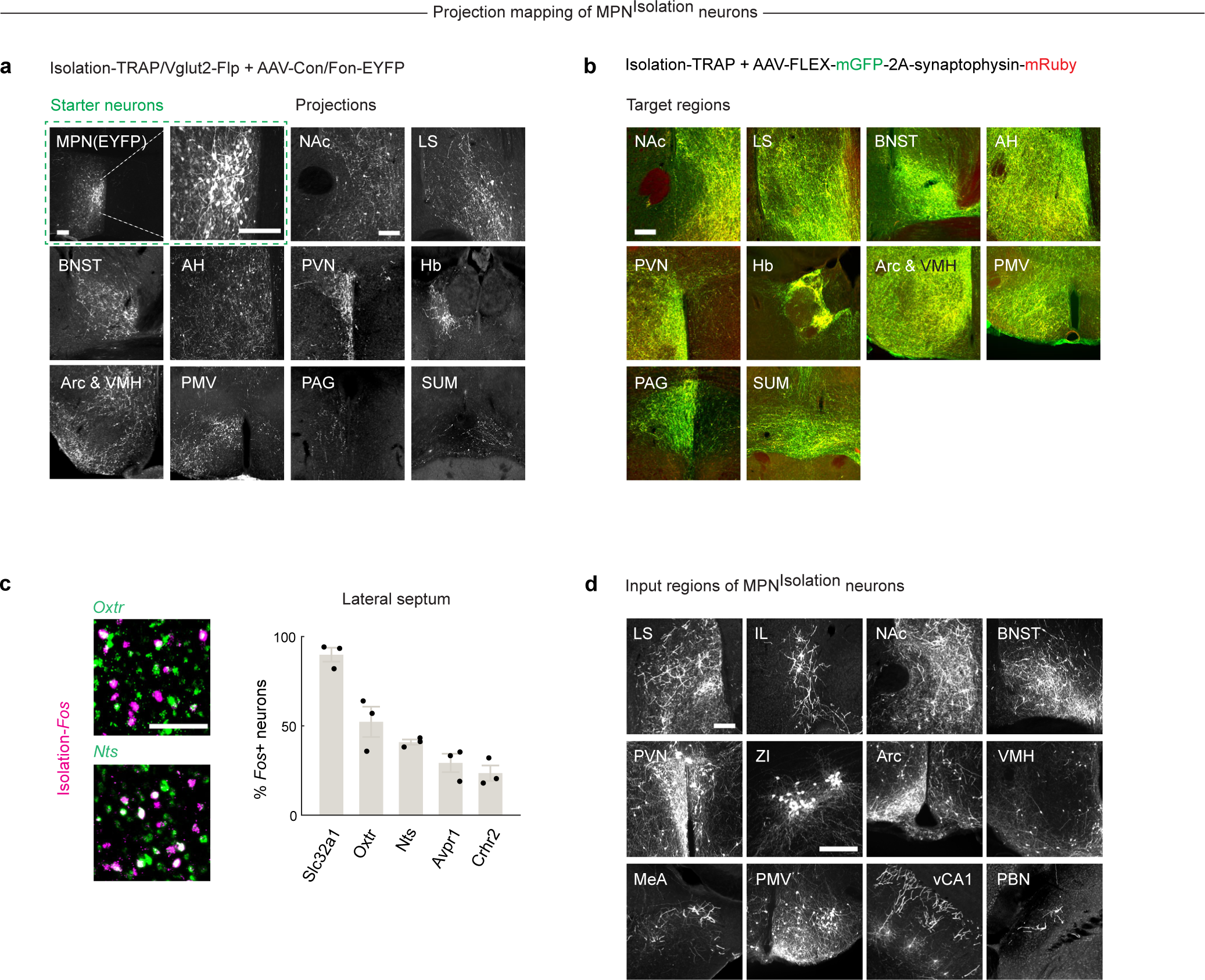
Circuit tracing from MPN^Isolation^ neurons. **a-b**, Representative images showing downstream targets of MPN^Isolation^ neurons using intersectional (Cre/Flp, **a**) and TRAP (**b**) methods, n=2-3. All scale bars, 200μm. **c,** Cell type identification of LS neurons activated during isolation n=3. Scale bar, 100μm. **c**, Representative images of input regions of MPN^Isolation^ neurons. Both scale bars, 200μm. Images without scale bars have same scale as LS image.

**Extended Data Figure 7.**
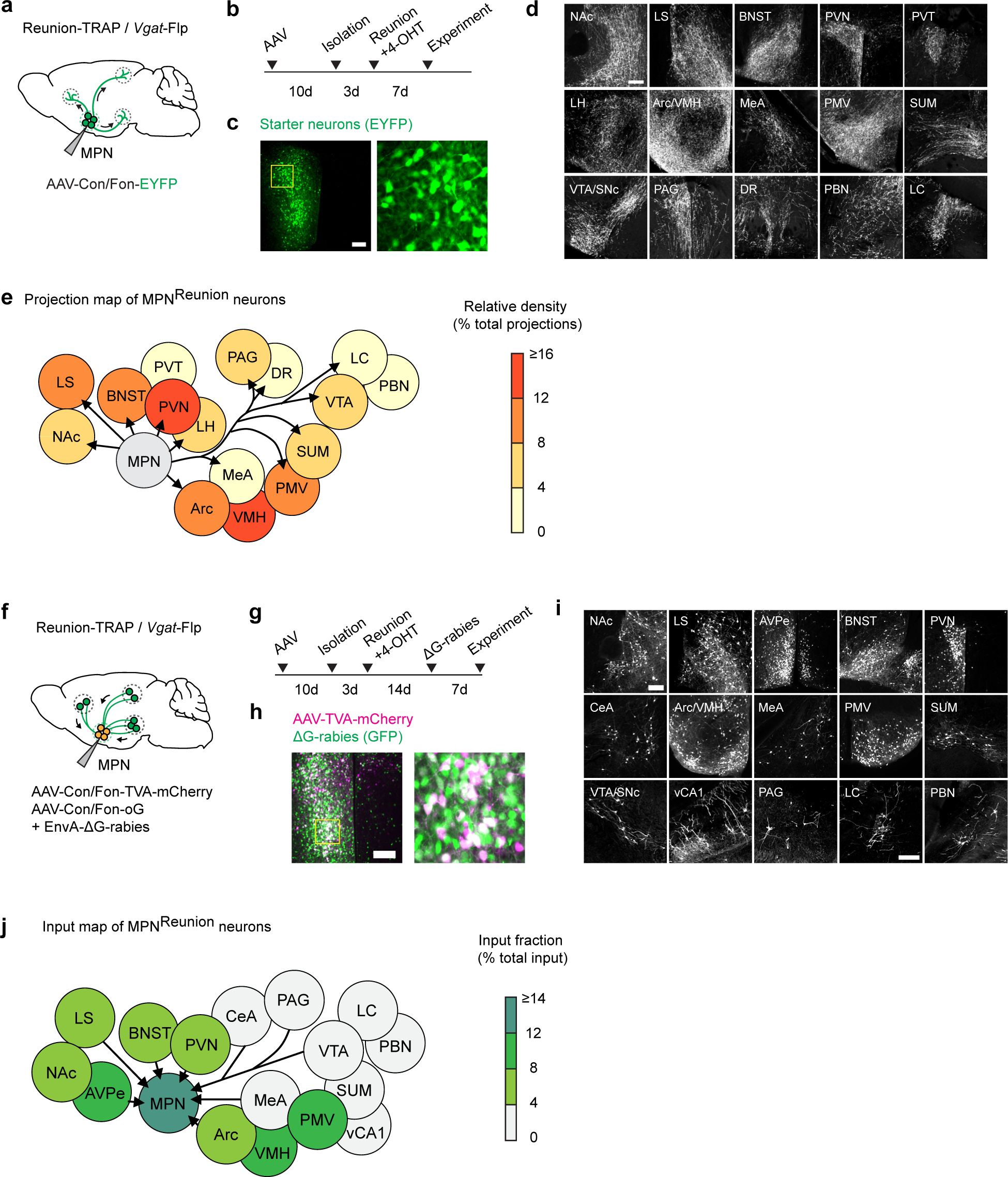
Circuit tracing from MPN^Reunion^ neurons. **a,b**, Viral tracing strategy to map target brain regions of MPN^Reunion^ neurons. **c**, Representative images of MPN^Reunion^ starter neurons. **d**, Representative images of projections from MPN^Reunion^ neurons. **e**, Schematic summary of projections from MPN^Reunion^ neurons and relative projection density. **f,g**, Viral tracing strategy to map input brain regions of MPN^Reunion^ neurons. **h**, Representative images of MPN^Reunion^ starter neurons in retrograde tracing. **i**, Representative images of input regions of MPN^Reunion^ neurons. **j**, Schematic summary of input brain regions of MPN^Reunion^ neurons and their relative input intensity. All scale bars, 200μm. Images without scale bars in **d** and **i** have same scales as in NAc images. Abbreviations: MPN, medial preoptic nucleus. NAc, nucleus accumbens. LS, lateral septum. AVPe, anteroventral periventricular nucleus. BNST, bed nucleus of the stria terminalis. PVN, paraventricular nucleus of hypothalamus. PVT, paraventricular thalamus. LH, lateral hypothalamus. CeA, central amygdala. Arc, arcuate nucleus. VMH, ventromedial hypothalamus. MeA, medial amygdala. PMV, ventral premammillary nucleus. SUM, supramammillary nucleus. VTA, ventral tegmental area. SNc, substantia nigra pars compacta. vCA1, ventral hippocampal CA1. PAG, periaqueductal gray. DR, dorsal raphe nucleus. PBN, parabrachial nucleus. LC, locus coeruleus.

**Extended Data Figure 8.**
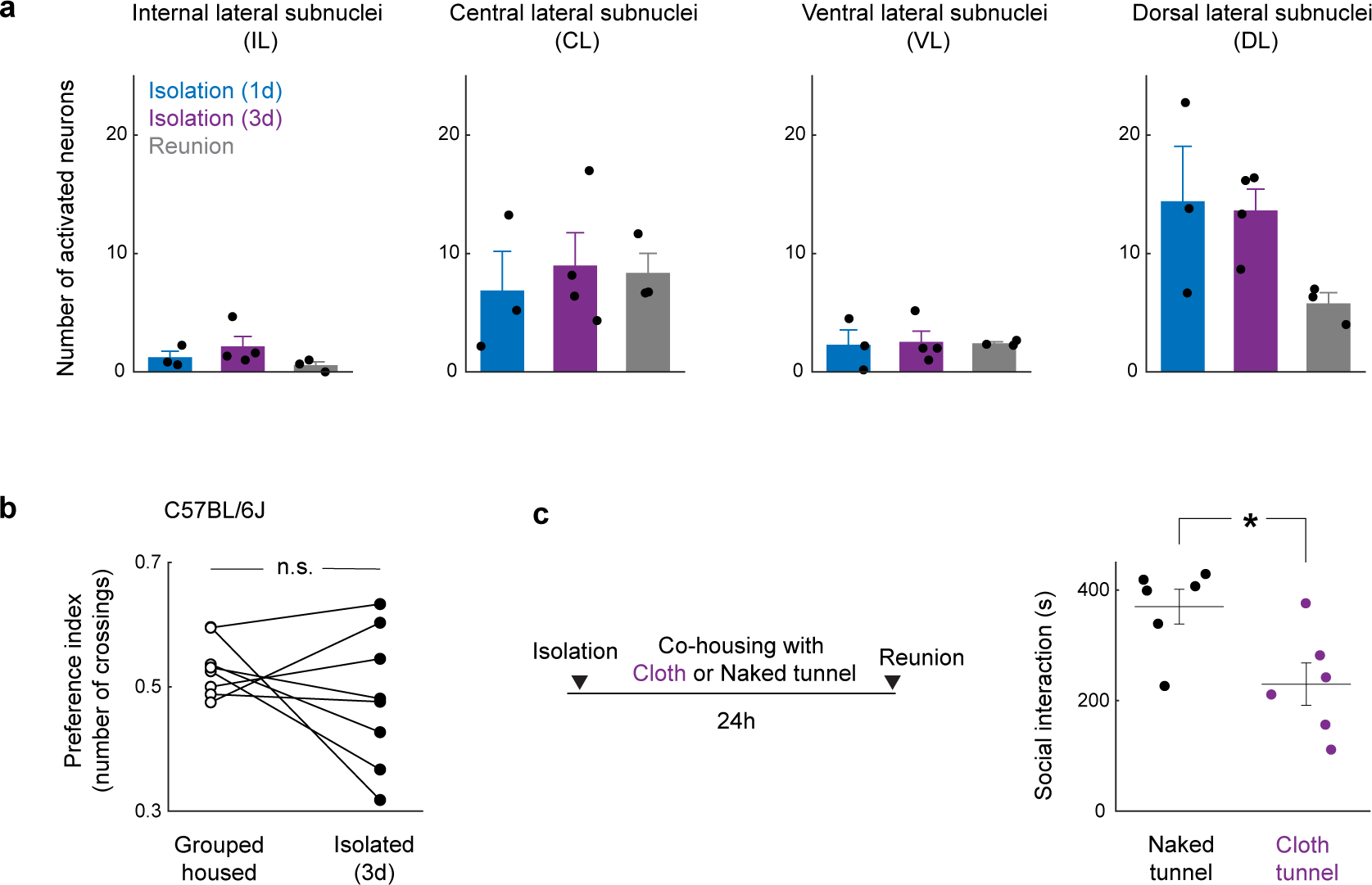
Modulation of social need by touch. **a**, Quantification of activated neurons in PBN subnuclei during social isolation and reunion, n=3-4 in each condition. Kruskal-Wallis test. **b**, Two-choice touch preference test in C57BL/6J mouse strain. Preference index is ratio of number of crossings through cloth tunnel over total number of crossings, n=7. Wilcoxon signed-rank tests; n.s., not significant. **c**, Behavioral effects of co-housing with cloth or naked tunnel during isolation on reunion social interaction, n=6. Mann– Whitney U test, *p<0.05. All error bars represent the mean ± s.e.m.

**Supplementary Video 1. Social interaction before and after social isolation, and examples of behavior modules in FVB/NJ mice**

Isolated mice display rebound social interaction upon reunion, while group housed mice show sated social motivation. These video clips illustrate typical behavior modules during social reunion, i.e. approaching, sniffing, crawling under, head-to-head contact and allogrooming.

**Supplementary Video 2. Microendoscopy calcium imaging of MPN neurons in FVB/NJ mice during social reunion.**

Examples of simultaneously imaged MPN neurons (right) showing distinct activity patterns during social reunion (left). The same color scheme was used as in Fig. 2d to indicate three unique neuronal clusters responding to social reunion.

**Supplementary Video 3. Comfort touch preference assay.**

Isolated but not group housed FVB/NJ mice show increased preference to cross a cloth-lined tunnel, rather than a naked tunnel.

## Methods

### Animals

Mice were maintained on a 12h:12h dark-light cycle with access to food and water ad libitum. The temperature was maintained at 22°C and the humidity between 30-70%. All experiments were performed in accordance with NIH guidelines and approved by the Harvard University Institutional Animal Care and Use Committee. Experiments were performed in adult female mice to avoid interfering behaviors such as aggression or mating seen in males after social isolation. The following mouse lines were obtained from the Jackson Laboratory: BALB/cJ (000651), DBA/2J (000671), C57BL/6J (000664), C3H/HeJ (000659), SWR/J (000689), FVB/NJ (001800), TRAP2 (also called Fos^2A-iCreERT2^, 030323), Ai9(RCL-tdT) (007909), Mc4r-2a-Cre (030759), Vglut2-IRES2-FlpO-D (030212), VGAT-2A-FlpO-D (029591), ROSA-DTA (009669). We obtained *Pde6b^rd1^* mutant mice in C57BL/6J background from Dr. Connie Cepko (Harvard Medical School), and Mrgprb4-Cre mice from Dr. Ishmail Abdus-Saboor (Columbia University). Trpc2 knockout mice were generated previously^39^ by Dr. Dulac. The following mouse lines were backcrossed to FVB/NJ mice for (N) generations before experiments: TRAP2/Ai9 (N>9, activity labeling experiments), Vglut2-Flp and VGAT-Flp (N≥5, neural tracing and optogenetics), Mc4r-Cre (N=3, calcium imaging and optogenetics), Mrgprb4/DTA and Trpc2 (N=1, behavioral assay). The sample sizes for experiments were chosen based on common practices in animal behavior experiments.

### Behavioral assays

All behavioral experiments were performed during the dark cycle of the animals in a room illuminated by infrared or red light. Mice were habituated in the room for 10-20 minutes before experiments. For stress tests, mice were acclimated to the testing environment for 1h before testing to reduce basal stress level.

#### Social isolation/reunion assay

Female sibling mice were group housed (≥3 mice) after weaning for at least one week before social isolation experiments. Before isolation, two group-housed cagemates were put together in a new cage with fresh bedding for 10 minutes to measure the baseline social interaction. One mouse was then isolated in their home cage or in a new cage for 5 days while the other mouse was kept in group. On the first, third and fifth day from the onset of isolation, the isolated mice were transiently reunited with the same group-housed cagemate in a new cage for 10 minutes. In each reunion session, the isolated mice were first put in the cage, and then its group-housed cagemate was introduced. All behaviors occurring during the reunion period were recorded using a multi-camera surveillance system (GeoVision GV-VMS software and GV-BX4700-3V cameras). Behavior videos were manually scored using the Observer XT 11 software (Noldus Information Technology) to identify typical social behavioral modules, including approaching (one mouse moves towards another), sniffing (the nose of one mouse comes close to or makes contact with another mouse’s body, usually the anogenital region), crawling under (one mouse crouches down, crawls underneath another mouse’s body and sometimes passes through), head-to-head contact (two mice approach each other and contact each other’s noses) and allogrooming (one mouse grooms another mouse, usually on the head, neck and back regions) (Fig. 1f and Extended Data Fig. 1e). Every single event of these behavioral modules was considered as a social interaction bout (Extended Data Fig. 1b, c), and the interval between the introduction of the second mouse and the initial social interaction bout was measured as behavioral latency (Extended Data Fig. 1a). The transition probability of one behavioral module occurring after a different module was calculated across all reunion sessions to quantify the specific motor sequences of social interaction during reunion (Extended Fig. 1d, transition matrix). The total duration of all the modules within one reunion session was summed up and referred to as social interaction (Fig. 1c, d). In order to demonstrate the satiation process during social rebound, we measured social interaction in separate time bins (2 minutes/bin) during reunion (Fig. 1j). Social interactions were also measured after 1, 3, 6 and 12 hours of isolation to reveal the emergence of social rebound after short periods of social isolation (Fig. 1i). Custom-made MATLAB codes and DeepLabCut software package were used to track frame-by-frame positions of two mice during social reunion and the social distances between two mice were calculated and averaged across frames during one reunion session (Fig. 1e). Because 1 and 3 days of social isolation are sufficient to trigger significant social rebound in FVB/NJ mice, we used these two time points in our typical isolation protocols in order to minimize increased stress induced by prolonged isolation.

#### Ultrasonic vocalization detection

Ultrasonic vocalizations (USVs) during social reunion were recorded in a sound isolation box using an ultrasonic microphone (Ultrasound Gate CM16/CMPA; Avisoft Bioacoustics) positioned 30 cm above the floor of the cage and converted into a digital format by an analog-to-digital (A/D) converter (Ultrasound Gate 116, Avisoft Bioacoustics) sampling at 500 kHz. The Avisoft Recorder software was used to control the recording and audio files were saved as 16 bit WAV format and later analyzed with DeepSqueak^51^, a deep learning-based system for detection and analysis of USVs. The built-in mouse call detecting neural network was used to identify USV syllables, and the detected syllables were manually reviewed to correct false labeling (see example syllables in Extended Data Fig. 1f). The corrected detection files were then exported as .mat files and analyzed with MATLAB. Custom code was used to extract the timing and duration of each syllable. The number of syllables were plotted as a function of different time courses (Fig. 1g and Extended Data Fig. 1g) to reveal the dynamic changes within and across reunions. As only one microphone was used for recording, we were not able to determine the source of USVs; however, since group-housed female mice emit few USVs and we detected USVs from isolated mice before cagemates were introduced, thus most of the recorded USVs during reunion were likely generated by the isolated mice. Correlation between USVs and social interaction initiated by the isolated mice were assessed over time (Fig. 1h and Extended Data Fig. 1h).

#### Non-social object interaction test

To test whether social isolation increases the motivation for general investigation behaviors, we tested behavior towards a non-social object in group housed or isolated conditions. Before isolation, single group-housed mice were presented with a 15ml centrifuge tube or a rubber toy mouse in a new cage for 10 minutes to measure baseline behavior. Mice were then isolated in their home cage or in a new cage for 3 days before a second test with the object. Behavior during tests was recorded and analyzed. Any contact or climbing behavior with the object was scored and added up as the time spent interacting with object (Fig. 1l).

#### Social interaction in different phases of estrous cycle

To test whether social rebound in female mice after social isolation is influenced by the estrous stage of the animal, we performed reunion assays after 3 days of social isolation and identified the estrous phase of each tested mouse. Time spent in social interaction was compared between mice in either estrus or diestrus phases (Fig. 1k). Vaginal smears were examined under the microscope, and specific estrous stages were characterized based on the morphology of vaginal epithelial cells as described previously^52^. In brief, we collected vaginal cells from female mice with 10µl of phosphate-buffered saline (PBS) and observed these samples under a light microscope with a 40x objective to characterize the morphology of cells. During the estrus phase, vaginal epithelial cells are cornified and appear large and flat, while during the diestrus phase, these cells are smaller with round shape^52^.

#### Stress tests

Mice’s stress levels were measured after different durations of social isolation using elevated plus maze (Fig. 1m) and open field (Fig. 1n) tests, and induced stress was performed by physical restraint (Fig. 1o). The elevated plus maze was on a pedestal 1m above the ground and consisted of two closed arms (30 x 5cm with 15-cm-high wall) and two open arms (30 x 5cm) arranged 90° from each other with a central platform (5 x 5cm), all made of black acrylic board. Mice were placed on the central platform and allowed to freely explore the maze for 5 minutes while the behavior was recorded. All behavior videos were manually scored with Noldus Observer software to measure the total time spent in open and closed arms. In the open field test, individual mice were placed in a 42 x 42 x 42cm arena composed of black acrylic board and allowed free exploration for 10 minutes. The behavior was recorded and analyzed to measure the total time spent in center (24 x 24cm) and peripheral zones. To induce physical restraint stress, individual mice were placed into 50ml conical tubes with ventilation holes for 1h and then reunited with one of their cagemates in a new cage to monitor the behaviors with the same settings as described for social reunion assays.

#### Social preference tests

To examine the preference of test mice for a group of mice versus a single mouse, we built a novel arena with three cubic chambers (25 x 25 x 25 cm) joined via a triangular central zone to allow for unbiased entry into either chamber (Fig. 1p). Each chamber contained an inverted wire cup. The cup was empty in the C0 chamber, contained one cagemate in the C1 chamber, and contained three cagemates in C3 chamber. The locations of the three types of chambers were assigned randomly across animals. Group-housed or isolated mice were first introduced into the central zone at the beginning of the test and allowed to freely explore the arena for 10 minutes. Behaviors were recorded and time spent in each chamber was scored manually. In the preference test between stranger versus familiar mice (Extended Data Fig. 1j), a standard three-chamber task was used in which the arena consisted of two choice chambers on two sides and a central zone that allowed for unbiased entry into either chamber. Each of the two choice chambers contained an inverted wire cup with either a familiar mouse (cagemate) or a stranger from the same strain. The tested mice (group-housed or isolated) were first introduced into the central zone at the beginning of the test and allowed to freely explore the arena for 10 minutes. Behavior was recorded and time spent in each chamber was scored. The same arena and experimental procedure were used in the strain preference test (Extended Data Fig. 1k), in which the two choice chambers contained mice from either the same or different strains.

#### Oxytocin receptor antagonist experiment

To examine the contribution of oxytocin to the regulation of social need during isolation, mice received i.p. injections of either saline or the oxytocin receptor antagonist (OTR-A), L-368,899 hydrochloride (5 mg/kg), twice during a 24h isolation as previously described^19^. The first injection was performed at the onset of isolation and the second injection 9h before the reunion assay (Fig. 3x). Behaviors during the reunion period were recorded and analyzed as described above.

#### Sensory modality screening

To investigate the contribution of different sensory modality to the emergence of social rebound after isolation, we designed a home-cage divider experiment in which a plastic divider with laser-cut thin slots/openings was placed in the diagonal of home cage to subdivide a group of 3 mice, such that one mouse was placed into one side and the other two together on the other side. Food, water and nesting materials were equally provided on both sides. The divider prevented mice from either side from physically interacting with each other but allowed exchange of auditory and olfactory information. After 3 days of separation, the singly divided mouse was reunited with one of the mice from the other side in a new cage for 10 minutes. The occurring behaviors were recorded and manually scored. The time spent in social interaction was compared to the social rebound after 3 days of isolation in the singly housed condition (Fig. 5a, b). To examine the potential contribution of pheromone sensing to social rebound, we assessed the behavior of *Trpc2*-/- mice that are genetically impaired in vomeronasal pheromone sensing. *Trpc2*-/- mice were first crossed to FVB/NJ strain for one generation, and the resulting *Trpc2*+/- mice were used to cross to each other in order to generate wildtype (*Trpc2*+/+) and mutant (*Trpc2*-/-) mice for isolation experiments. We measured the rebound social interaction after 3 days of isolation and analyzed the satiation process in wildtype and mutant mice (Fig. 5c) with the same methods described in *Social isolation/reunion assay*.

#### Gentle touch preference test

To test the preference of gentle touch before and after social isolation, we designed a free choice task for mice interact with either a naked plastic tunnel (10-cm long, provided by Harvard Biological Research Infrastructure) or a tunnel lined inside with a single layer of soft plush towel (bought from Amazon) (referred to as cloth tunnel). All materials were autoclaved before use and replaced between animals to avoid odor contamination. Group-housed or 1-day isolated mice were placed in a new cage with one naked tunnel and one cloth tunnel. During the test, mice were allowed to freely explore and go through either tunnel for 15 minutes. Behaviors were recorded, and the crossing events were scored. The touch preference index was measured as the number of crossings though the cloth tunnel divided by the total number of crossings through both naked and cloth tunnels (Fig. 5g and Extended Data Fig. 8b).

#### Touch manipulation assays

To acutely reduce tactile sensitivity during social reunion, isolated mice received an i.p. injection of isoguvacine^40^ (20 mg/kg), a peripherally restricted GABA_A_R agonist, 60 minutes before social reunion. Phosphate-buffered saline (PBS) was injected in a different batch of isolated mice, which served as a control group. The injected mice were subjected to a social reunion assay 1 hour after injection, and the behaviors were recorded and manually scored (Fig. 5d). To examine the contribution of tactile sensation to the emergence of social need, we genetically ablated a population of mechanosensory neurons marked by Mrgprb4+ that are thought to mediate pleasant touch in mice^42,43^. We first separately crossed Mrgprb4-Cre (B4-Cre) and Cre-dependent diphtheria toxin subunit A (DTA) mouse lines (in C57BL/6J background) to FVB/NJ strain for one generation and then crossed the resulting F1s (i.e. F1(B4/FVB) and F1(DTA/FVB)) to each other to have B4-neuron-lesioned mice (B4/DTA). We measured the social interaction of B4/DTA mice towards a wildtype cagemate after 1 or 3 days of isolation and compared these results to the social rebound measured in wildtype mice from FVB/NJ x C57BL/6J cross (Fig. 5f). In the acute touch rescue experiments (Fig. 5h), faux-fur-lined tunnels were used to provide comfort touch stimulation as an enhanced version of cloth tunnel described in *Gentle touch preference test*. Mice were habituated to the faux-fur-lined tunnels by continuously going through two of these tunnels that were alternatingly connected by experimenter before social isolation. Mice were then isolated for 24 hours and reunited with one cagemate both before and after gentle touch stimulation. Specifically, a 5-minute reunion assay was first performed to measure the baseline social interaction, and then the mice went across faux-fur-lined tunnels 30 times the same way as in habituation. 30 times of tunnel crossings typically took ∼10 minutes for both cloth tunnel and naked control tunnel. A second reunion assay was then conducted to measure the change of social motivation compared to the first reunion (Fig. 5h). Naked plastic tunnels were used in another batch of animals as a negative control. In the chronic touch rescue experiment (Extended Data Fig. 8c), isolated mice were co-housed with either a cloth tunnel or a naked tunnel described in *Gentle touch preference test* for 24 hours. Then a social reunion assay was carried out to measure and compare social interaction after different co-housing conditions.

### Microendoscopy calcium imaging

To examine real-time neuronal activity with single cell resolution in the medial preoptic nucleus (MPN) of freely behaving animals, we performed microendoscopy calcium imaging in FVB/NJ mice (n=3) and Mc4r-Cre/FVB mice (n=3). The Mc4r-Cre mouse line was backcrossed for at least 3 generations to FVB/NJ strain before experiments. All imaging experiments were performed during the dark cycle of the animals in a room illuminated by dim red light.

#### Virus injection and GRIN lens implantation

For pan-neuronal activity imaging 400nl of AAV1-Syn-GCaMP6s (Addgene #100843-AAV1) was injected unilaterally into the MPN of FVB/NJ mice, at (AP 0, ML 0.3 and DV −4.8, Paxinos and Franklin atlas). To image MPN*^Mc4r^*^+^ neurons, 400nl of AAV1-Syn-Flex-GCaMP6s (Addgene #100845-AAV1) was injected unilaterally into the MPN of Mc4r-Cre/FVB mice using the same coordinates. Since the brain anatomy of FVB/NJ strain slightly differs from the Paxinos and Franklin brain atlas, we adjusted the AP coordinate 0.4-0.5mm towards the rostral side to target the MPN in FVB/NJ mice. 30 minutes after viral injection, a 25-gauge blunt needle (SAI infusion technologies, VWR # 89134-146) was slowly inserted (1mm/5min) into the brain targeting (AP 0, ML 0.3 and DV −4.7, Paxinos and Franklin atlas) to create a tract. The needle then was slowly withdrawn, and a GRIN lens (Inscopix, 0.6 x 7.3 mm) was slowly inserted (1mm/10min) into the tract formed by the needle and targeted at 50µm below the end of the needle tract. The lens assembly was secured on the skull with dental cement (Parkell) and a titanium headplate was attached at the base of the lens assembly with dental cement to restrict the animal’s head for attaching to the microendoscope. The positions of all implanted GRIN lenses were assessed through post hoc histology and only the imaging data from correctly targeted lens were used for further analysis. Mice were individually housed after surgery for 1 week and then co-housed with 1 or 2 cagemate(s) for another 3-4 weeks to allow for the expression of GCaMP and clearing of the imaging window.

#### Calcium imaging and behavioral assays

Optimal imaging settings were determined on the first day of the imaging experiments. The implanted mouse was transiently restrained and the microendoscope (nVista3, Inscopix) was attached onto the lens assembly on the head of the mouse. The focusing plane of the microendoscope was carefully adjusted over the entire working distance to choose an imaging plane with most neurons and sharp image. The field of view was cropped to the region encompassing the fluorescent neurons. The illumination power (∼10% of the max power) and the sensor gain (∼10-20% of the maximum gain) were chosen to have a strong dF/F signal, but not saturated. These imaging settings were saved for each animal and subsequently used for the same animal across sessions. This allowed us to reliably image from the same field of view in different experiments and register the same neurons across sessions. We used inbuilt acquisition software from Inscopix to acquire images at 10Hz. Before formal data acquisition, mice were habituated to the recording setup and environment 2-3 times, and for each recording session, mice were habituated for 10-15 minutes with microendoscope before starting the behavioral assays. An Arduino microcontroller is programmed to send TTL signals to synchronize the data acquisition of calcium signals (DAQ, Inscopix), animal behavior (BFS-U3-31S4M-C, FLIR Blackfly S camera) and ultrasonic vocalization (Avisoft Bioacoustics). In the social reunion assay (Fig. 2b, c, k, l, m, n and Extended Data Fig. 2a, b, d), implanted mice were isolated for 3 days before imaging session. On the day of imaging, the mouse was first habituated in a new cage with fresh bedding for 10-15 minutes. Then the imaging session started and continued for 15 minutes. Initially, there was a 5-minute baseline period during which the mouse was kept alone. Following this, a cagemate was introduced into the cage, and social interaction between the mice was allowed for 5 minutes. Subsequently, the cagemate was removed, and the mouse was kept alone again for another 5 minutes. In the group housed condition (Fig. 2o), implanted mouse stayed with cagemates before imaging for at least 48 hours and during the imaging session, there was a 5-minute baseline during which the mouse was kept alone, followed by a 5-minute reunion with one cagemate, and then this cagemate was removed for another 5 minutes. Imaging during tail suspension (Fig. 2p) was conducted when the mouse was manually suspended by its tail for up to 2 minutes in order to induce a state of stress. Imaging during eating behavior (Fig. 2q) was performed when the mouse was fasted for 12h in group housing, and it was allowed for freely eating of food pellets in a new cage for 10 minutes during which the imaging was performed. To monitor the neuronal activity during the initial phase of social isolation, we performed imaging when the mouse was initially isolated from its social group for 20 minutes in a new cage (Extended Data Fig. 2e). We provided food and hydrogel in the cage and kept the mouse isolated for another 6 hours with the microendoscope connected. Next, we performed a 10-minute imaging with 5 minutes of baseline and 5 minutes of reunion. In the tunnel crossing session (Fig. 5i), the implanted mouse was isolated for 3 days before imaging. During the imaging session, there was a 5-minute baseline followed by 15 minutes of free tunnel crossing during which two faux-fur-lined tunnels (10cm long, 7.5cm in diameter, the fur ∼1.5 cm long) were introduced into the cage to maximize crossing behaviors. The top of the tunnel was removed creating a slot (4cm) to allow the implanted mice to run through with the attached wire. The tunnels were subsequently removed from the cage and the mouse was kept alone for 30 minutes. Finally, we performed a 10-minute imaging with 5 minutes of baseline and 5 minutes of reunion.

#### Image processing and calcium signal extraction

The images acquired were processed in two steps. In the first step, we processed the raw image data using the Inscopix data processing software (IDPS 1.8.0.3519). The images were imported in the proprietary Inscopix format on to IDPS. The images were spatially down sampled by a factor of 4 in order to reduce the file sizes for subsequent steps without losing quality, followed by a spatial bandpass filter in the frequency band of 0.005 to 0.5/pixel. The images were next subjected to motion correction using IDPS and the resulting images were saved as a tiff image stack. The second step was performed using standard MATLAB scripts from the CNMFe^53^ database (https://github.com/zhoupc/CNMF_E). Calcium traces were extracted and deconvolved using CNMFe pipeline with the following parameters: patch_par=[2,2], gSig = 3, gSiz = 13, ring_radius = 9, min_corr = 0.8, min_pnr = 8, deconvolution: foopsi with the ar1 model^53^. The spatial and temporal components of every extracted unit were carefully inspected manually (SNR, PNR, size, motion artifacts, decay kinetics, etc.) and outliers (obvious deviations from the normal distribution) were discarded. Cells with elongated and thin shape were removed based on their shape.

#### Single neuron and population activity analysis

All analysis based on extracted calcium traces were performed using custom made MATLAB scripts. To visualize and categorize the pan-neuronal calcium activity patterns during social reunion assay (Fig. 2c, d and Extended Data Fig. 2a, b), the extracted calcium traces were z-scored across the whole imaging session, and the neurons with enhanced or inhibited activity during social reunion period were separated by comparing the mean activity during 5min baseline and 5min reunion. The reunion-activated neurons were then further divided into two clusters with distinctive activation patterns (one cluster with transient activation at the onset of reunion and the other cluster with persistent activation during reunion period) using k-mean clustering (k=2). We visualized the resulting 3 clusters (i.e. reunion-inhibited, reunion-activated-transient and reunion-activated-persistent) in heatmaps in single-neuron level (Fig. 2c and Extended Data Fig. 2a) and averaged activity traces across all neurons from the same cluster (Fig. 2d and Extended Data Fig. 2b). To visualize the population activity during social reunion assay, we performed principal component analysis (PCA) across the simultaneously recorded neurons and projected the activity onto the first three PCs to track population trajectory across time (bin size=5s, Fig. 2e). To calculate the population trajectory distance from baseline as a measure of state change, the Euclidian distance between each point in the trajectory and the mean of the baseline (-100 to 0s) was calculated for each time bin (1s). The same procedure was repeated 100 times by randomly selecting 50% of the neurons to calculate the standard deviation (Fig. 2f).

In the activity analysis of MPN^Mc4r+^ neurons, significantly modulated neurons were identified based on their responses during reunion. Reunion-activated neurons are defined as activated at least two standard deviations above the mean of baseline (300s before reunion). Reunion-inhibited neurons are identified as significantly reduced activity during reunion period (300s after reunion) compared to baseline (300s before reunion, Mann–Whitney U test, bin size=1s). We visualized the activity of all significantly activated or inhibited Mc4r+ neurons from three mice (Fig. 2k, l and Extended Data Fig. 2d) aligned at reunion time and plotted their corresponding activity during re-isolation period aligned at the time point when the partner mouse was removed from the cage. Calcium traces from 10 example neurons (2 reunion-activated, 7 reunion-inhibited and 1 not significantly modulated) were selected and plotted in Fig. 3l. We plotted the averaged activity of the reunion-inhibited neurons during reunion (Fig. 2m) and re-isolation (Fig. 2n). We identified these reunion-inhibited neurons in other behavioral conditions by alignment of fields of view in different recording sessions. The spatial location of each neuron is binarized and a proportion of overlap is calculated for every pair of neurons across the sessions. If the proportion of overlap is higher than a set threshold (>70%), the neurons were considered to be the same. We plotted the averaged activity of reunion-inhibited neurons during social interaction in group housed animals (Fig. 2o), tail suspension (Fig. 2p) and eating (Fig. 2q). The significance of modulation was determined by testing whether the activity of neurons was above or below 95% confidence interval of baseline (time bin = 1s). To examine the activity of reunion-inhibited neurons during the initial phase of social isolation (Extended Data Fig. 2e) and soft cloth tunnel crossing (Fig. 5i), the raw image data were concatenated with later recordings during social reunion, and calcium traces were extracted from the concatenated video using CNMFe as described above. Reunion modulated neurons were identified and their activity during the early recording period was plotted. In the activity heatmap of the initial phase of isolation (Extended Data Fig. 2e), neurons were arranged according to their onset of activation after isolation (from early to late responses). The curve below represents the average activity across neurons in each time bin (30s). In the analysis of cloth-tunnel crossing experiments, two touch-activated and two touch-inhibited neurons were selected, and the average activity of multiple crossings were plotted that is aligned at the onset of each tunnel crossing (Fig. 5j). To assess the co-modulation of neuronal activity between tunnel crossing and social reunion, the activity changes during tunnel crossing (mean(crossing, 30s) – mean(baseline,10s)) and social reunion (mean(reunion, 300s –mean(baseline, 300s)) were plotted for each neuron in Fig. 5k.

### Cell type identification

#### TRAP induction and activity labelling

To specifically label neurons activated during social isolation, we took the advantage of the TRAP2/Ai9 mouse line^17^ that enables the labeling of activated neurons over a long time window (3-6 hours), which better integrates neuronal activity representing a persistent state compared to *Fos in situ* hybridization. The TRAP2/Ai9 line was backcrossed to FVB/NJ strain for over 9 generations before use. The isolation-activated neurons were ‘TRAPed’ in the following steps. 10mg of 4-hydroxytamoxifen (4-OHT, Sigma Aldrich #H6278-10mg) was added with 500μl ethanol and shaken at room temperature until the powder was completely dissolved (20mg/ml). This solution could be frozen at −20°C for future use within 1 month. The resolved 4-OHT solution was then mixed with corn oil in 1:2 volume ratio and vortexed to fully mixed to extract the 4-OHT in the oil. The resulting solution was vacuum centrifuged for 1-1.5 hour until the upper layer of ethanol was evaporated. The final 10mg/ml 4-OHT solution was intraperitoneally injected into mice immediately after preparation at a dose of 50mg/kg. To label (‘TRAP’) the cells activated during social isolation, mice were isolated for 3 days and injected with the 4-OHT solution in the dark phase of the third day and immediately returned to the same isolation cage for at least 24 hours to allow tamoxifen fully metabolized. After at least 7 days, the mice were sacrificed for in situ hybridization experiments. To assess the activity of PBN during isolation, we performed TRAP inductions after 1 or 3 days of isolation in different cohorts of animals. To label the neurons that are activated during social reunion, FVB/NJ mice were isolated for 3 days and reunited with one of its cagemate. 30-40 minutes after the onset of the reunion, mice were sacrificed for in situ hybridization experiments. The reunited mice were kept with cagemate from reunion assay until sacrifice to prevent the activation of isolation-related neurons. To assess the activity of PBN during reunion, 4-OHT was injected 1h after reunion and the reunited mice were kept in group for at least 24 hours to allow tamoxifen fully metabolized.

#### *In situ* hybridization

RNAscope V2 kit (Advanced Cell Diagnostics, ACD) was used to perform double-label and triple-label fluorescence in situ hybridization (FISH) according to the manufacturer’s instructions. Probes for *tdTomato* and *Fos* were used to visualize activated neurons labelled with TRAP method or acute behavioral assays, respectively. Probes for marker genes of specific cell types were selected from previous single-cell RNA sequencing and functional studies. All probes were made by ACD. Animals were sacrificed after specific behavioral assays and the brains were dissected. Freshly frozen brains were sectioned using a cryostat at 16 μm and stored at −80 °C. On the day of FISH experiment, slides were thawed and fixed in 4% paraformaldehyde (PFA) for 15 min followed by dehydration in 50%, 75% and 100% ethanol at room temperature. Tissue samples were processed using 3% hydrogen peroxide (VWR) for 10 minutes and permeabilized for 25 minutes using Protease IV (ACD). For each RNAscope experiment, C2 and C3 probes were diluted in C1 probe solution (1:50), heated to 40°C for 10 minutes and applied to slides which were placed in ACD HybEZ II oven at 40°C for 2 hours. Tissue samples were then processed as suggested by the RNAscope V2 protocol (ACD). Slides were imaged at 10x on an Axioscan 7 using Zen Blue 3.5 software (Zeiss). The number of cells marked by specific and overlapping genes was measured using QuPath 0.3.2.

### Neural circuit tracing

#### Anterograde tracing

Anterograde tracing experiments were performed in TRAP2(Fos-CreER)/Vglut2-Flp mice and TRAP2/Vgat-Flp mice both in FVB/NJ background. TRAP2, Vglut2-Flp and Vgat-Flp lines were separately backcrossed to FVB/NJ strain for more than 5 generations and the offspring from these lines were crossed to generate TRAP2/Vglut2-Flp/FVB mice for tracing experiments of MPN^Isolation^ neurons and TRAP2/Vgat-Flp/FVB mice for tracing experiments of MPN^Reunion^ neurons. All surgeries were performed under aseptic conditions in animals anaesthetized with 100mg/kg ketamine and 10mg/kg xylazine via intraperitoneal injection. Using a programmable nano-injector (Nanoject III, Drummond), 150-200nl of Cre- and Flp-dependent virus AAV8-hSyn-Con/Fon-EYFP (Addgene #55650-AAV8) was unilaterally injected into the MPN (AP 0, ML 0.3 and DV −5, Paxinos and Franklin atlas). We adjusted the AP coordinate 0.4-0.5mm towards the rostral side to match the anatomy of FVB/NJ brain. After surgery, injected mice were singly housed to recover for 1 week and then put together with former cagemates for another week before TRAP induction (see details in *TRAP induction and activity labelling*). To visualize the projections of MPN^Isolation^ neurons, TRAP2/Vglut2-Flp/FVB mice were isolated for 3 days and injected with 4-OHT (Fig. 3j,k). To visualize the projections of MPN^Reunion^ neurons, TRAP2/Vglut2-Flp/FVB mice were isolated for 3 days and injected with 4-OHT one hour after reunion (Extended Fig. 7a, b). After 2 weeks of viral fluorophore expression, animals were perfused transcardially with phosphate-buffered saline (PBS) followed by 4% paraformaldehyde (PFA) in PBS. Brains were dissected and post-fixed in 4% PFA overnight. After embedding in 4% low-melting point agarose (Promega, #V2111) in PBS, 50-μm coronal sections were cut through the whole brain on a vibratome (Leica) and mounted on slides (VWR, 48311-703) with DAPI-containing mounting medium (Vector Laboratories, H-1200). The brain sections were imaged at 10x magnification using AxioScan 7 and Zen Blue 3.5 software (Zeiss). For quantification of projection density, the average pixel intensity in a target region containing EYFP signals was calculated, and the background was subtracted (Zen Blue software, Zeiss). Because injections were unilateral and no labelling was observed in most cases contralaterally, the equivalent region on the contralateral hemisphere was chosen for background subtraction; in cases where contralateral EYFP were present, an adjacent unlabeled region was chosen. The relative density value for each projection region was calculated as the ratio between background-corrected intensities in each region divided by the sum across all the target regions (Fig. 3l and Extended Data Fig. 7e).

#### Monosynaptic retrograde tracing

Monosynaptic retrograde tracing experiments were performed with an intersectional strategy in TRAP2/Vglut2-Flp/FVB mice and TRAP2/Vgat-Flp/FVB mice (see details in *Anterograde tracing*). We first unilaterally injected 150-200nl of a 1:1 mixture of two Cre- and Flp-dependent viruses, AAV8-nEF-Con/Fon-TVA-mCherry (Stanford GVVC-AAV-197) and AAV8-Ef1a-Con/Fon-oG (Stanford GVVC-AAV-198), into the MPN with the same coordinates as in *Anterograde tracing*. The injected mice were singly housed for 1 week and reunited with former cagemates for another week before TRAP induction (see details in *TRAP induction and activity labelling*). TRAP2/Vglut2-Flp/FVB mice were isolated for 3 days and injected with 4-OHT to allow virus expression in MPN^Isolation^ neurons (Fig. 4a, b); TRAP2/Vgat-Flp/FVB mice were isolated for 3 days and injected with 4-OHT one hour after reunion to enable virus expression in MPN^Reunion^ neurons (Extended Data Fig. 7f,g). Two weeks later, 200nl of G-deleted rabies virus (EnvA-ΔG-rabies-eGFP, Janelia Viral Tools Facility) was injected into the MPN. Seven days later, mice were euthanized, and the brains were dissected, sectioned and imaged with the same procedure in *Anterograde tracing.* Relative input strength was quantified as follows. First the representative sections of input regions were selected and GFP+ cells (presynaptic cells) were counted. The local presynaptic cells within the MPN were estimated by counting GFP+ and mCherry-neurons. The relative input density was calculated as the ratio between number of presynaptic cells in each input region divided by the sum across all calculated regions in each brain (Fig. 4d and Extended Data Fig. 7j). To identify the input cell types of MPN^Isolation^ neurons, a different cohort of mice were processed for *in situ* hybridization (see details in *In situ hybridization*). Probes for *GFP*, marker genes or immediate early genes were used to examine the presynaptic cell types (Fig. 4e).

### Optogenetics

#### Virus injection and fiber implantation

TRAP2/Vglut2-Flp/FVB mice and TRAP2/Vgat-Flp/FVB mice were bilaterally injected with 200nl of AAV8-hSyn Con/Fon-hChR2(H134R)-EYFP (for activation; Addgene #55645) or AAV8-nEF-Con/Fon-iC++-EYFP (for inhibition; Addgene # 137155) into the MPN (AP 0, ML ±0.3 and DV −5, Paxinos and Franklin atlas) and in the same surgery a dual fiber-optic cannula (200/250-0.66_GS0.6/0.8_FLT, Doric Lenses) was implanted 200µm above the injection site for MPN cell body manipulation or above the Arc (AP -1.6, ML ±0.3 and DV −5.3) or the Hb (AP -1.6, ML ±0.3 and DV −2.2) for projecting axon terminal manipulation. To manipulate MPN Mc4r+ neurons, 200nl of AAV-EF1a-DIO-hChR2(H134R)-EYFP (activation, UNC Vector Core) or AAV-EF1a-DIO-iC++-EYFP (inhibition, UNC Vector Core) was bilaterally injected into the MPN and the dual fiber-optic cannula was implanted. Mice were recovered for 2 weeks before Isolation-TRAP and Reunion-TRAP induction (see details in *TRAP induction and activity labelling*). Mice were tested 3–5 weeks after TRAP induction to allow for efficient expression of ChR2 or iC++.

#### Optogenetic manipulations

On testing days, the implanted optic fibers were attached through a patch cord (SBP(2)_200/230/900-0.57_FCM-GS0.6/0.8, Doric Lenses) and a rotary joint (FRJ_1x1_FC-FC, Doric Lenses) to a 460-nm blue LED module (Prizmatix) for optogenetic activation or inhibition. An Arduino microcontroller is programmed to send TTL signals to LED module to control the stimulation patterns. Pilot experiments were conducted to test and determine the proper ranges of LED power in different manipulation experiments. After connected to the patch cord, mice were transferred to a new cage with the same setup in social reunion assay and allowed to habituate for 5-10 minutes. To activate Vglut2+/Isolation-TRAPed neurons (Fig. 3d-f) or Mc4r+ neurons (Extended Data Fig. 5b), LED was on for 1s (20Hz, 10ms pulses, 6-8mW at patch cord tip) and off for 3s, repeatedly; To activate Vgat+/Reunion-TRAPed neurons (Fig. 4h,j), LED was on for 40s (20Hz, 20ms pulses, 6-8mW) and off for 20s, repeatedly; In the real-time place preference/avoidance tests (Fig. 3g, 4i and Extended Data Fig. 5c), LED was turned on (20Hz, 6-8mW) during the period when mice entered the LED-on chamber that is randomly assigned in each session. To inhibit Vglut2+/Isolation-TRAPed neurons (Fig. 3h,i), Mc4r+ neurons (Extended Data Fig. 5d) or Vgat+/Reunion-TRAPed neurons (Fig. 4k), LED was on for 3 minute (constant on, 3-4mW) and off for 20s, repeatedly. In the social interaction/reunion tests and three-chamber social preference tests (Fig. 3d, e, h, i; Fig. 4h, j, k and Extended Data Fig. 5b, d), patterned LED was applied for 10 minutes and the behaviors during the entire period were recorded and analyzed as LED-on session performance. In LED-off sessions, the same cohorts of animals were tested without LED stimulation. LED-off sessions were usually conducted before LED-on sessions to avoid possible conditioning effects after stimulation. In the pre-stimulation experiment (Fig. 3f), patterned LED was applied for 10 minutes when the animal was kept alone before a 10-minute reunion with LED off. The behaviors during the reunion period were recorded and analyzed. The real-time place preference/avoidance tests lasted for 10 minutes, and the patterned LED was applied when mice entered the LED-on chamber. For axon terminal manipulation experiments (Fig. 3p-w), we used the same protocols as in soma stimulation experiments described above. When testing the stimulation effects on food intake, mice were fasted for 24 hours before experiments. Two food pellets were placed on the two sides of the cage and animal contact with one pellet triggered LED on for 10s while contact with the other pellet did not trigger stimulation treated as off controls. Weight reduction of each pellet at the end of a 10-minute test was measured as the amount of food intake (Fig. 3q,u).

### Fiber photometry

#### Virus injection and fiber implantation

All surgeries were performed under aseptic conditions with animals anesthetized with isoflurane (1–2% at 0.5–1.0 L/min). Analgesia was administered pre-(buprenorphine, 0.1 mg/kg, i.p.) and post-operatively (ketoprofen, 5 mg/kg, i.p.). We used the following coordinates to target virus injection and fiber implant for the nucleus accumbens core (NAc): (AP 1.78, ML 1.0 and DV −3.6 from dura, Paxinos and Franklin atlas). To express dopamine sensor GRAB_DA2m_^36^, we unilaterally injected 300nl of mixed (3:1) virus solution: AAV9-Syn-GRABDA2m (Vigene Bioscience) and AAV5-CAG-tdTomato (UNC Vector Core) into the NAc. Virus injection lasted around 10 minutes, after which the injection pipette was slowly removed over the course of several minutes to prevent damage to the tissue. We then implanted an optic fiber (400 µm diameter, SMA-SMC, Doric Lenses) into the virus injection site. We first slowly lowered an optical fiber into the NAc, attached it to the skull with UV-curing epoxy (NOA81, Thorlabs), and then a layer of rapid-curing epoxy to attach the optical fiber more firmly to the underlying glue. After 15 minutes, we applied a black dental adhesive (Ortho-Jet, Lang Dental) and waited for another 15 minutes for it to dry. The implanted mice were singly housed for 1 week to recover and co-housed with previous cagemates for another week before photometry recording.

#### Fiber photometry recording

Photometry recording was performed as previously reported^54,55^. Before recording, we connected a magnetic patch cord (400µm diameter, NA 0.48, 3m long, SMA-SMC, Doric Lenses) to the optical fiber implanted on the head of the animal and the animal was allowed to habituate in a new cage for 10-15 minutes. Once the recording started, the patch cord simultaneously delivered excitation light at different wavelength (473nm, Laserglow Technologies; 561 nm, Opto Engine LLC) and collected fluorescence emissions from dopamine sensor and tdTomato (used for motion correction). The emitted light was then filtered using a 493/574nm beam splitter (Semrock), followed by a 500 ± 20nm (Chroma) and 661 ± 20nm (Semrock) bandpass filter, and collected by a photodetector (FDS10 X 10 silicone photodiode, Thorlabs) connected to a current preamplifier (SR570, Stanford Research Systems). This preamplifier outputs a voltage signal which was collected by a data acquisition board (NIDAQ, National Instruments) and custom software written in Labview (National Instruments). Lasers were turned on at least 30 minutes prior to recording to allow them to stabilize. Before each recording session, laser power and amplifier settings were individually adjusted for each mouse. The photometry recording and behavior video acquisition were synchronized using a common TTL input to trigger infrared light pulses (once every 10 seconds) that were recorded in the behavior videos. The implanted mice were isolated for 3 days before recording. The recording session lasted for 10 minutes, including a 5-minute baseline period where the animal was kept alone, followed by a 5-minute social reunion period where a previous cagemate was introduced.

#### Fiber photometry data analysis

The tdTomato signal (red) was subtracted from dopamine sensor signal (green) to correct the motion artifacts. The corrected signal was then z-scored using the mean and standard deviation from a 30-second ‘baseline period’ before reunion (Fig. 4n, upper: example mouse). Individual traces from different mice were aligned at the reunion time point and averaged across animals (Fig. 4n, lower). The behaviors during recording sessions were manually scored and synchronized with photometry signals by common TTL pulses (see *Fiber photometry recording* above).

## Statistics and reproducibility

Data were processed and analyzed using MATLAB and GraphPad Prism 9. The sample sizes were chosen based on common practices in animal behavior experiments. Individual data points were plotted wherever possible. Error bars and shaded areas in the graphs indicate the mean ± s.e.m. unless otherwise noted. All data were analyzed with two-tailed non-parametric tests unless otherwise noted. In the experiments with paired samples, we used the Wilcoxon matched-pairs signed-rank test or Friedman test. In the experiments with non-paired samples, we used the Mann–Whitney U test or Kruskal-Wallis test. P values were corrected for multiple comparisons when necessary. No statistical significance is indicted by n.s., and the significance was indicated by *P < 0.05, **P < 0.01, ***P < 0.001. Statistical details are given in the respective figure legends. All behavioral, imaging, in situ hybridization, optogenetics and tracing experiments were replicated in multiple batches of animals independently with similar results. Experiments were randomized whenever possible. Experimenters were blind to the mouse identify in Mrgprb4+ neuron lesion experiments.

## Data availability

All data that support the findings of this study are either present in figures and extended data or available from the Dulac lab GitHub database (https://github.com/DulacLabHarvard/SocialNeed) or from the corresponding author upon request.

## Code availability

Custom codes written for behavioral and imaging data analysis are available at the Dulac Lab GitHub database (https://github.com/DulacLabHarvard/SocialNeed) or from the corresponding author upon request.

## Notes

### Competing Interest Statement

The authors have declared no competing interest.

